# A Distinct Oxygenase Family Catalyzing Methyllanthionine Sulfoxide Formation in Anti-phage Class I Lanthipeptides

**DOI:** 10.64898/2026.08.26.747419

**Authors:** Jingqi Chen, Lingyang Zhu, Wilfred van der Donk

## Abstract

Lanthipeptides are one of the largest classes of ribosomally synthesized and post-translationally modified peptides (RiPPs). The *coi* biosynthetic gene cluster (BGC) from *Streptomyces coelicolor* A3(2) encodes a canonical class I lanthipeptide dehydratase (CoiB) and cyclase (CoiC), a bifunctional enzyme (CoiS_A_) with an *O*-methyltransferase (MT) and glutamyl lyase (GL) domain, and a protein of unknown function (CoiH). The product of the *coi* BGC was recently shown to impart anti-phage activity, but its structure is still unresolved. Previous work investigated the regioselectivity of the GL domains in CoiB and CoiS_A_ and the stereochemistry of the cyclized precursor peptide, but the function of CoiH was not addressed. In this study, co-expression of the peptide CoiA1 with CoiBCS_A_H resulted in a +16 Da addition to the cyclized peptide compared to when CoiH was omitted. LC-MS/MS analysis indicated that this modification occurred in the first of three thioether rings. A combination of site-directed mutagenesis, comparison of linear and cyclized peptide substrates, hydrogen peroxide (H₂O₂) treatment, and collision-induced dissociation (CID) mass spectrometric analysis suggested that the sulfur atom in the first methyllanthionine was oxidized to a sulfoxide by CoiH. This hypothesis was confirmed by NMR analysis. CoiH represents a previously uncharacterized oxygenase family catalyzing sulfoxide formation. Structure prediction tools suggest a new enzyme fold without obvious metal or cofactor binding sites, although the possibility that CoiH utilizes cofactors or other cellular components for catalysis cannot be excluded.

## INTRODUCTION

Lanthipeptides represent one of the largest classes of ribosomally synthesized and posttranslationally modified peptides (RiPPs).^1,2^ They are characterized by thioether-containing lanthionine (Lan) and/or methyllanthionine (MeLan) residues. The biosynthesis of lanthipeptides involves the dehydration of Ser/Thr residues followed by subsequent cyclization of Cys residues onto the dehydroalanine (Dha) or dehydrobutyrine (Dhb) to generate (Me)Lan (Figure S1).^1,3^ Depending on the biosynthetic enzymes, lanthipeptides are classified into five classes (I to V). Conventional class I lanthipeptide biosynthesis features a dehydratase LanB that activates the side-chain hydroxyl groups of Ser and Thr residues by glutamylation using Glu-tRNA (Figure S1). In a second active site, the glutamyl group is then eliminated by a glutamyl lyase domain (GL).^4^ A cyclase LanC catalyzes the Michael-type addition of Cys thiols onto the dehydroamino acids to form (Me)Lan.^5^

Previously, the *coi* BGC from *Streptomyces coelicolor* A3(2) was shown to encode three precursor peptides (CoiA1-3), and a canonical class I dehydratase CoiB and cyclase CoiC.^6^ In addition, the *coi* BGC encodes a bifunctional enzyme CoiS_A_ composed of a C-terminal *O*-methyltransferase and an N-terminal second GL domain, and a previously uncharacterized protein CoiH (Figure 1A). Both GL domains in CoiB and CoiS_A_ catalyze glutamate elimination; however, they were shown to generate two different Dhb isomers, with the GL in CoiB generating *Z*-Dhb and the GL in CoiS_A_ forming *E*-Dhb.^7,8^ The enzyme OlvS_A_, which contains a similar methyltransferase domain as CoiS_A_ does not contain a GL domain, methylates a conserved Asp residue resulting in formation of an isoAsp.^9^ In the absence of CoiH, the ring pattern of the CoiA1 product formed by CoiBCS_A_ features an N-terminal LL-MeLan ring and overlapping C-terminal DL- and D-*allo*-L-MeLan rings (Figure 1B).^7^ The overall structure of CoiA1 modified by CoiBCS_A_ (hereafter called CoiA1BCS_A_) is the most complex of any lanthipeptide characterized to date in terms of the stereochemical configuration of MeLan residues.^7^ Although the role of CoiB and CoiS_A_ in the biosynthesis of CoiA1BCS_A_ and the resulting stereochemistry have been well characterized, the function of CoiH and the structure of the final product are still unclear. Bioinformatic analysis shows widespread distribution of *coiH*-like genes. Their consistent association with conserved class I lanthipeptide BGCs suggests an important biological role. Recently, a study reported that the *coi* and related BGCs are ubiquitous in Actinobacteria and that their products are involved in defense against phage.^10^ Thus, the structure of the lanthipeptides that confer this activity termed lanthivirins^10^ is of high interest.

**Figure 1.**
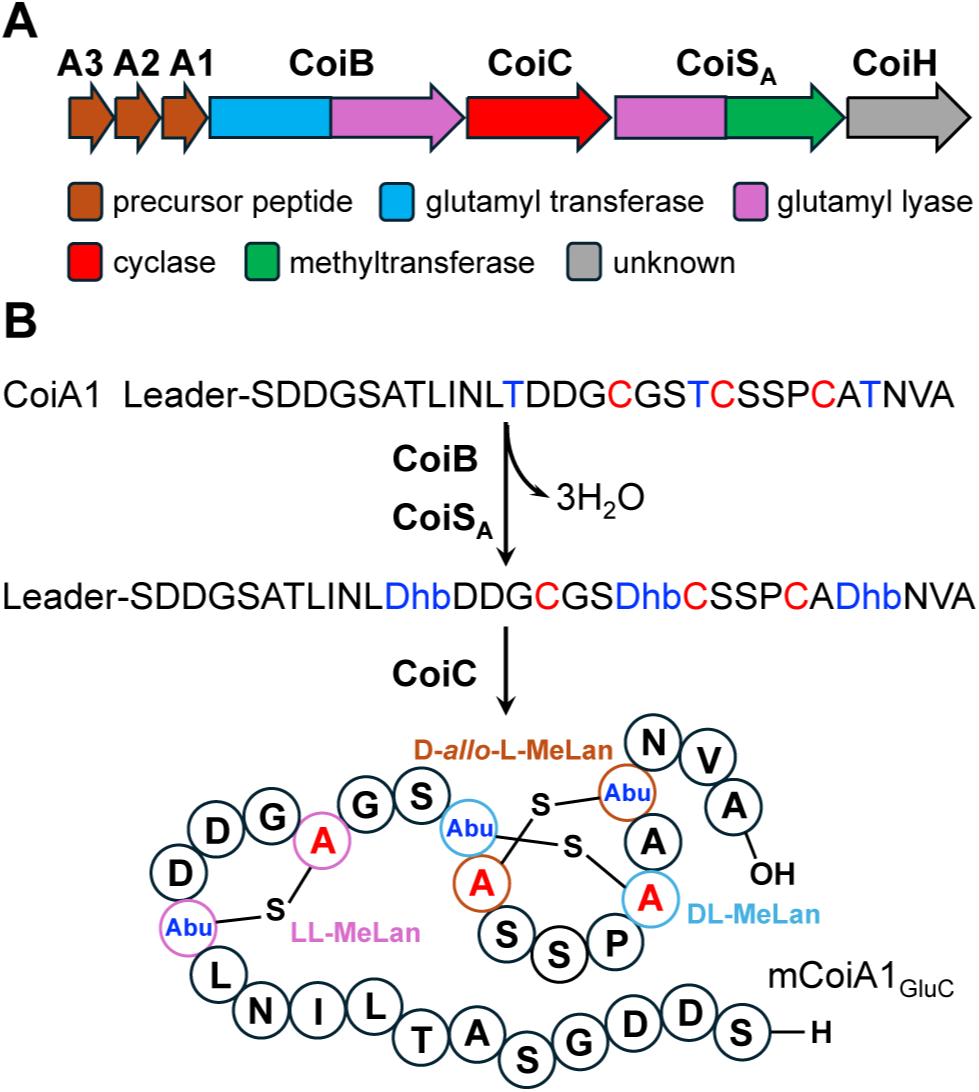
(A) The *coi* class I lanthipeptide BGC from *Streptomyces coelicolor* A3(2). (B) Biosynthesis of CoiA1BCS_A_ and the structure of the endoproteinase GluC-digested product. For convenience, the N-terminal part generated by GluC digestion is regarded as the leader peptide, but the exact site of leader peptide removal, if any, is not known. The sequence of the peptide that is indicated here as the leader peptide is MNANTIKGQAHSPAATAGGDAFDLDISVLE. mCoiA1, modified CoiA1. Dhb, dehydrobutyrine. Abu, 2-aminobutyric acid. For simplicity, the process is shown as complete dehydration by CoiB/CoiS_A_ followed by cyclization by CoiC, but CoiB/CoiS_A_ and CoiC could catalyze alternating dehydrations and cyclizations.

## RESULTS

### Coexpression of CoiA1 with CoiBCS_A_ and CoiH

Because a putative protease could not be identified within the *coi* BGC or related BGCs, we initially sought to characterize the final product by heterologous expression of the entire *coi* BGC in *Streptomyces lividans* TK-24 and *Streptomyces albus* J1074. Despite significant efforts, the mature product could not be detected or isolated from either host, consistent with similar efforts in a recent report on the anti-phage activity of the *coi* BGC.^10^ We then co-expressed CoiH with His_6_-SUMO-CoiA1, CoiB, CoiC, and CoiS_A_ in *E. coli* (heretoforth designated SUMO-CoiA1(BCS_A_H)). Upon purification by immobilized metal affinity chromatography (IMAC) and removal of the His_6_-SUMO tag using TEV protease and further digestion by endoproteinase GluC, MALDI-TOF MS revealed a +16 Da shift from the three-fold dehydrated peptide produced when CoiH was absent (Figure 2A). LC-MS/MS of the endoproteinase GluC-digested peptide with this 16 Da addition suggested that the modification of +16 Da occurred in the first thioether ring, but the exact position of the modification could not be resolved (Figure 2C). Aside from the addition of 16 Da in the first ring, the fragmentation pattern was the same as that obtained without CoiH expression (Figure 2B), suggesting that the overall ring pattern was not changed.

**Figure 2.**
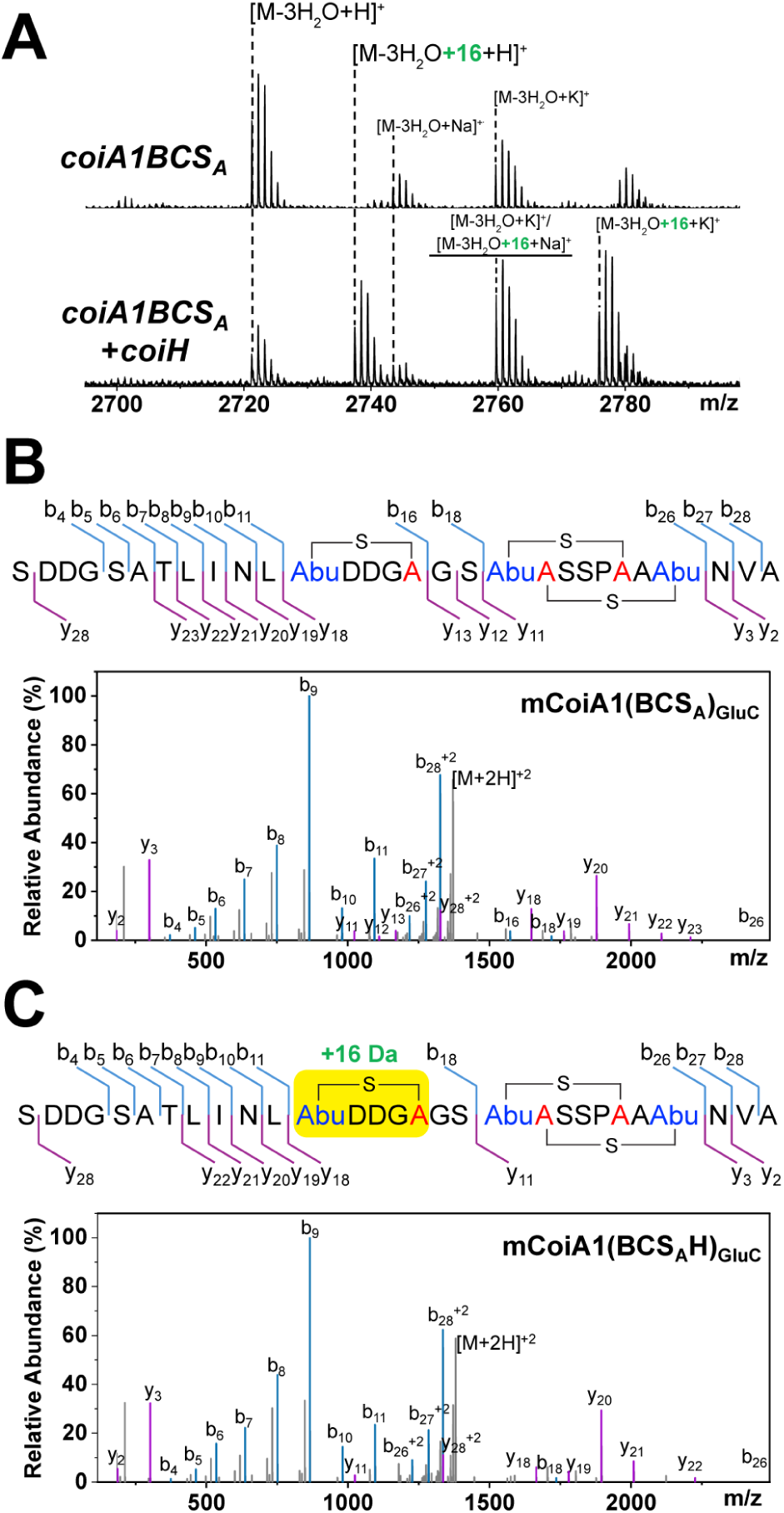
(A) Matrix-assisted laser desorption ionization time-of-flight (MALDI-TOF) MS analysis of coexpression experiments in *E. coli*. Coexpressed genes/proteins are listed on each panel. An N-terminal SUMO tag was fused to CoiA1 to enhance peptide solubility during heterologous expression. After purification, the fusion protein was first digested with TEV protease to remove the SUMO tag, followed by digestion with endoproteinase GluC. (B) ESI LC-MS/MS analysis of the GluC-digested mCoiA1(BCS_A_). mCoiA1(BCS_A_) denotes CoiA1 modified by CoiBCS_A_. (C) ESI LC-MS/MS analysis of the CoiBCS_A_H-modified CoiA1 (mCoiA1(BCS_A_H)) after GluC digestion. Fragmentation results are consistent with a 16 Da addition on the N-terminal MeLan (yellow-highlighted region) and two C-terminal overlapping MeLan. For calculated and observed m/z values of molecular ions and fragment ions, see Table S1.

### CoiH does not catalyze hydroxylation of conserved Asp residues

The highly conserved double Asp motif in the first ring area (Figure S2) was considered a potential modification target given the frequent hydroxylation of these residues in RiPPs.^2,11–16^ To investigate their possible involvement, either one or both Asp residues were mutated to Ala residues. Constructs encoding the SUMO-CoiA1 variants D43A, D44A, and D43A/D44A were generated by site-directed mutagenesis, and these peptides were co-expressed with CoiB, CoiC, CoiS_A_, and CoiH in *E. coli* followed by isolation of the product by Ni-affinity chromatography (Figure 3A). MALDI-TOF MS analysis showed that although the efficiency varied, all three CoiA1 variants underwent not only 3-fold dehydration but also the +16 Da modification (Figure 3B). Hence, the conserved Asp residues in the first ring are not the CoiH-modification sites. Because oxidation of a thioether ring to yield a cyclic sulfoxide was previously reported in the biosynthesis of the lantibiotic actagardine (Figure S3A),^17^ CoiH could perform a similar function to the corresponding monooxygenase GarO in actagardine biosynthesis (Figure S3B).^18,19^ To verify this hypothesis, we first determined whether CoiH could modify the linear dehydrated precursor peptide. A truncated CoiA1 analog (CoiA1_1-48_) in which the core peptide product would only contain the N-terminal methyllanthionine ring sequence was designed (Figure 3A). His_6_-SUMO-CoiA1_1-48_ was co-expressed with CoiB, CoiS_A_ and CoiH in *E. coli*, and a negative control was included in which His_6_-SUMO-CoiA1_1-48_ was co-expressed with only CoiB and CoiS_A_. MALDI-TOF MS analysis did not reveal any differences in mass of the modified product from both experiments, which suggests that dehydrated but uncyclized CoiA1_1-48_ is not a substrate for CoiH. In contrast, when CoiC was also co-expressed, the +16 Da mass shift was observed in the product peptide (designated mCoiA1_1-48_(BCS_A_H)) (Figure 3C). We therefore conclude that the formation of the first methyllanthionine ring is a prerequisite for the subsequent +16 Da modification by CoiH. This observation provided further support to the hypothesis that CoiH catalyzes methyllanthionine sulfoxidation.

**Figure 3.**
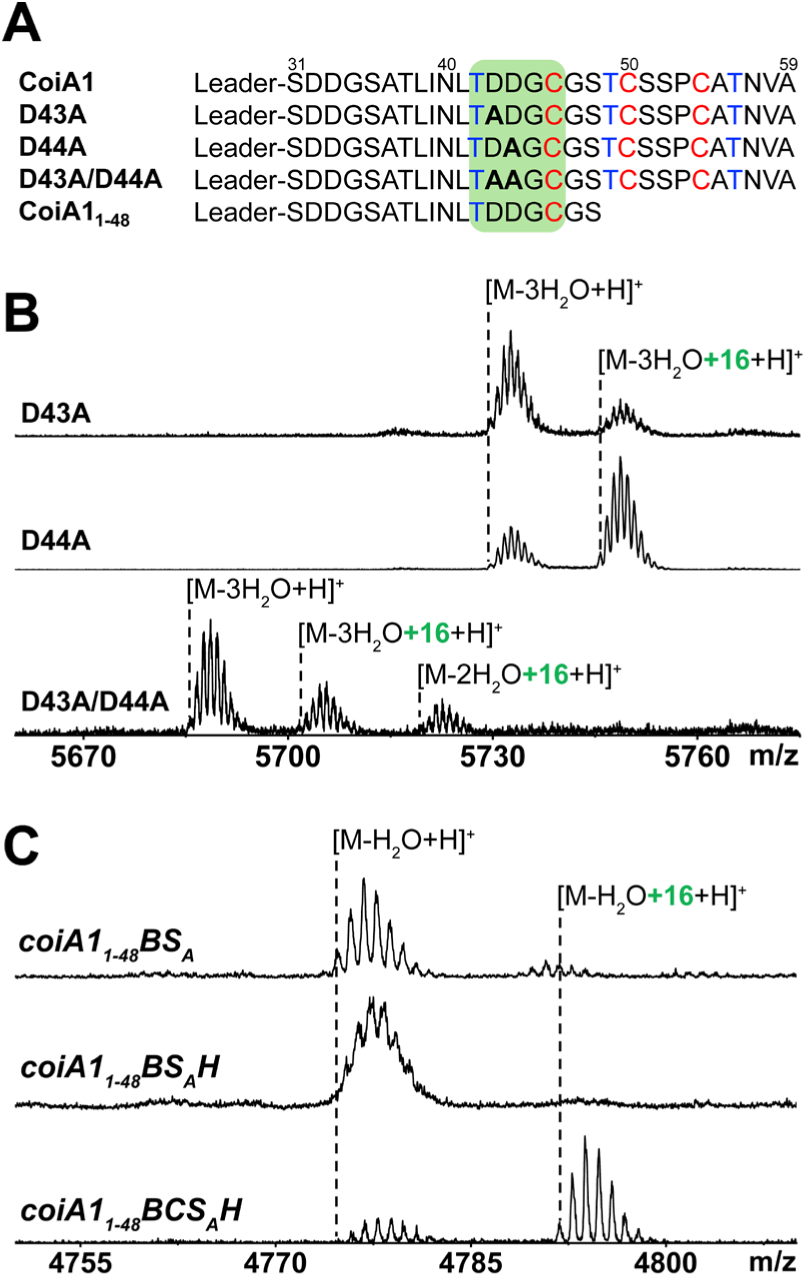
(A) Sequences of full-length CoiA1 and its variants. The first thioether ring sequence is highlighted by the green shaded region. (B) MALDI-TOF MS analysis of CoiA1 variants D43A, D44A, and D43A/D44A coexpressed with CoiB, CoiC, CoiS_A_ and CoiH. (C) MALDI-TOF MS analysis of the truncated peptide CoiA1_1-48_ coexpressed with different combinations of enzymes from the *coi* BGC. In panels B and C, the purified product was cleaved by TEV protease to remove the His_6_-SUMO tag. For calculated and observed m/z values of molecular ions, see Table S2.

### CoiH catalyzes oxidation of the methyllanthionine A-ring

The modification site and type of modification introduced by CoiH can be unambiguously determined by NMR analysis. However, purification of CoiA1 or CoiA1_1-48_ modified by CoiBCS_A_ and digested with GluC proved challenging, and the resulting peptide did not have sufficient purity nor yield for NMR characterization. We therefore employed two independent indirect approaches to verify the CoiH-mediated modification. Hydrogen peroxide (H_2_O_2_) is known to oxidize sulfur-containing amino acid residues, including methionine,^20^ and thioether-containing residues such as lanthionine and methyllanthionine, yielding the corresponding sulfoxides. TEV protease-digested SUMO-mCoiA1_1-48_ that had been co-expressed with CoiBCS_A_H (mCoiA1_1-48_(BCS_A_H)_TEV_) was treated with H_2_O_2_ at 37 ℃ for 4 h. The reaction was subsequently quenched by the addition of sodium pyruvate.^21^ MALDI-TOF MS analysis showed that the two enzymatically modified forms of the CoiA1_1-48_ peptide (M – H_2_O and M – H_2_O + 16) were converted to a single product, corresponding to mass increases of +32 Da and +16 Da compared to their respective peptides before H_2_O_2_ treatment (Figure 4A). Thus, one new H_2_O_2_-mediated oxidation had occurred for the CoiA1_1-48_ peptide with one dehydration and the +16 Da modification. LC-MS/MS data support that this additional oxidation caused by H_2_O_2_ was localized to the Met residue in the leader peptide part (Figure 4B). To provide even more clear data, Ala36 in CoiA1_1-48_ was substituted with Lys, allowing the generation of a shorter core peptide lacking the leader peptide (and its Met) through LysC protease digestion. Incubation with H_2_O_2_ resulted in a single oxidation of the CoiBCS_A_-modified peptide (mCoiA1_1-48_-A36K(BCS_A_)_LysC_) (Figure 4C and Figure S4); however, no further oxidation was observed upon H_2_O_2_ treatment for CoiH-modified product (mCoiA1_1-48_-A36K(BCS_A_H)_LysC_) (Figure 4D and Figure S5). These data support that CoiH oxidized the thioether linkage; if the oxidation occurred anywhere else, products with three oxidations should have been observed upon H_2_O_2_ treatment (Met, MeLan, and CoiH oxidation site) for mCoiA1_1-48_(BCS_A_H)_TEV_, or products with one oxidation should have been detected upon H_2_O_2_ treatment for mCoiA1_1-48_-A36K(BCS_A_H)_LysC_.

**Figure 4.**
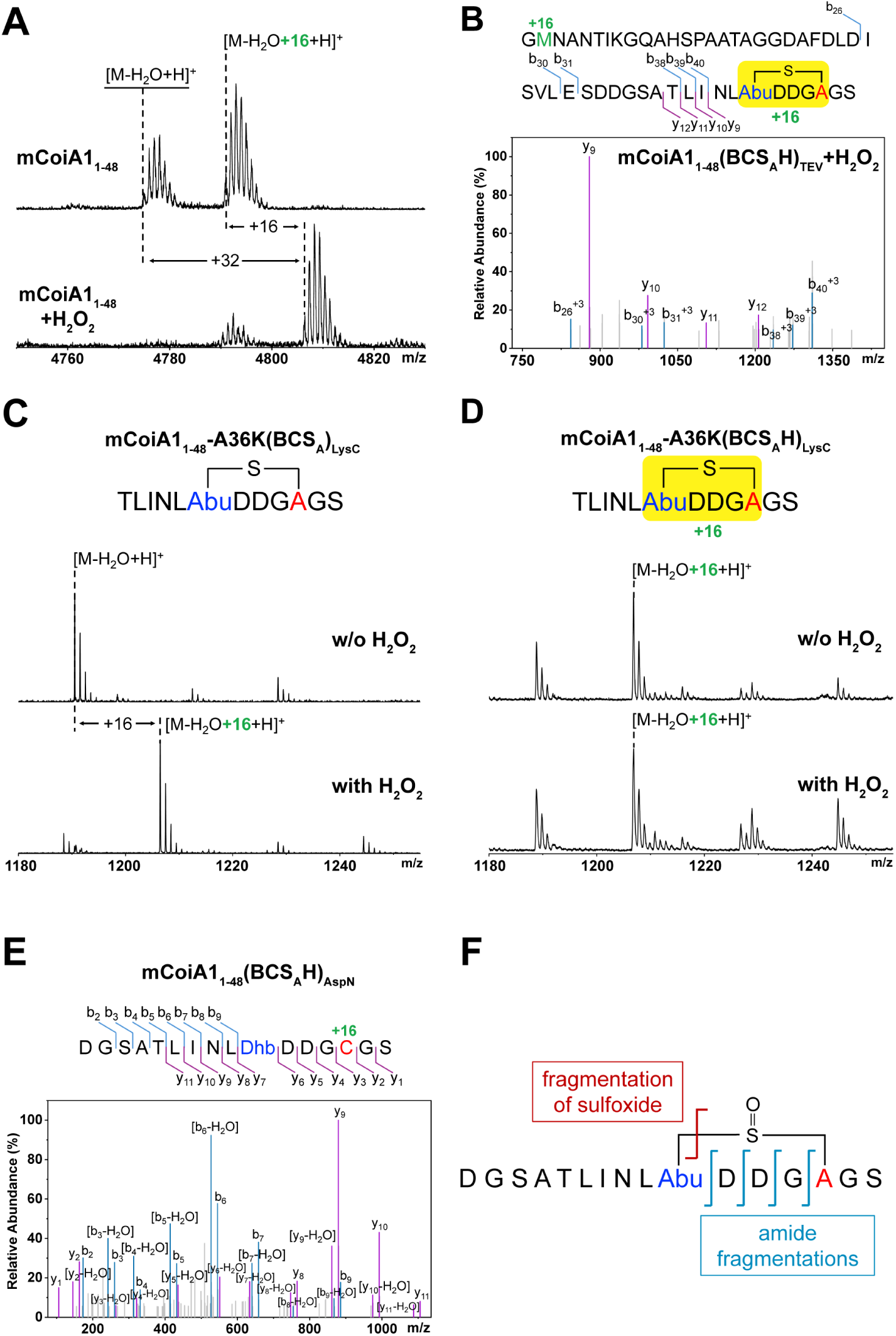
(A) Oxidation of mCoiA1_1-48_(BCS_A_H)_TEV_ by H_2_O_2_. The peptides were analyzed by MALDI-TOF MS. (B) LC-MS/MS analysis of the H_2_O_2_-oxidized mCoiA1_1-48_(BCS_A_H)_TEV_. In panels A and B, prior to H_2_O_2_ treatment, the SUMO-tagged mCoiA1_1-48_(BCS_A_H) was digested with TEV protease, leaving an additional Gly residue at the N-terminus. For LC-MS/MS analysis, the fragmentation pattern shows one +16 Da adduct on Met2, and one +16 Da adduct in the C-terminal MeLan region. (C) MALDI-TOF MS analysis after oxidation of mCoiA1_1-48_-A36K(BCS_A_)_LysC_ by H_2_O_2_. The corresponding LC-MS/MS data are shown in Figure S4-S5. (D) MALDI-TOF MS analysis for H_2_O_2_ treatment of mCoiA1_1-48_-A36K(BCS_A_H)_LysC_. (E) LC-MS/MS analysis of AspN-digested mCoiA1_1-48_(BCS_A_H) at elevated collision energies, locating the +16 Da to the Cys of the MeLan. (F) CID-induced sulfoxide and amide bond cleavage according to the LC-MS/MS data in panel E. For calculated and observed m/z values of molecular ions and fragment ions, see Table S3.

Thioether linkages are typically highly stable and exhibit limited fragmentation under conventional collision-induced dissociation (CID) conditions. This characteristic has been extensively utilized in RiPP studies to determine thioether ring formation and cyclization patterns.^22^ A recent study demonstrated that conversion of thioether bridges to sulfoxides significantly facilitates CID-mediated bond cleavage.^23^ According to the enhanced fragmentation properties, a mass spectrometry-based sequencing method for thioether-containing macrocyclic peptides was developed.^23^ Applying the same principle, the AspN-digested mCoiA1_1-48_(BCS_A_H) was subjected to LC-MS/MS analysis at elevated collision energies. Cleavage of amide bonds within the ring and the putative sulfoxide-containing thioether linkage was observed (Figure 4E). Because sulfoxide formation is known to markedly enhance CID-mediated cleavage of thioether bonds, these fragmentation patterns provide indirect additional evidence that mCoiA1_1-48_(BCS_A_H) contains an oxidized thioether residue (Figure 4F).

### NMR analysis confirms formation of a sulfoxide by CoiH

Collectively, all the evidence strongly supports that CoiH catalyzes an oxidation reaction of the N-terminal methyllanthionine in CoiA1 to form a sulfoxide. To confirm this conclusion, we revisited our attempts to obtain a peptide sample suitable for NMR characterization. Following large scale (12 L) co-expression of His_6_-SUMO-CoiA1_1-48_-A36K with CoiBCS_A_H in *E. coli*, purification of the mutant peptide by IMAC, and digestion by LysC, a 12-amino-acid peptide was obtained and purified by HPLC for NMR analysis (Figure 5A-B). The peptide structure was elucidated using 1D and 2D NMR experiments, including ^1^H, ^1^H-^1^H TOCSY, ^1^H-^1^H NOESY, and ^1^H-^13^C HSQC (Figure 5C-D, and Figures S6-S9). The ^1^H and ^13^C chemical shift assignments of the peptide in 90% H_2_O/10% D_2_O and in 100% D_2_O at 25 °C are summarized in Table S5. Both the ^1^H and ^13^C chemical shift values at the β-position of former Thr6 (for residue numbering, see Figure 5B) showed significant deviations from those of a typical Thr residue, suggesting its involvement in methyllanthionine formation. This conclusion was supported by the NOESY spectrum, which shows cross-peaks between the β-protons of former Thr6 (3.74 ppm) and former Cys10 (3.44 and 3.37 ppm), as well as between γ-protons of former Thr6 (1.16 ppm) and the β-protons of former Cys10, supporting a linkage between these two residues (Figure 5C). In addition, the ^13^C chemical shifts of the β-carbons are 60.6 ppm for the former Thr6 and 48.3 ppm for the former Cys10 (Figure 5D). These values are substantially downfield relative to the β-carbon chemical shifts of the methyllanthionine linkage in mCoiA1_1-48_-A36K(BCS_A_)_LysC_ (i.e. not converted to the sulfoxide by CoiH) (42.1 ppm for the former Thr and 29.8 ppm for the former Cys) (Figure 5F),^24^ indicating the influence of a strongly electron-withdrawing substituent. These perturbations in chemical shifts are in the same direction as those reported for the sulfoxide methyllanthionine in actagardine (Cβ derived from Thr at 54.4 ppm, and Cβ derived from Cys at 51.8 ppm).^17^ Moreover, the Cγ of the former Thr (methyl group) is shifted in the opposite direction (upfield) to 9.2 ppm (6 ppm in actagardine) from 18.2 ppm upon CoiH modification of the peptide (Figure 5D and F), which is consistent with the known β effect of sulfoxides.^25^

**Figure 5.**
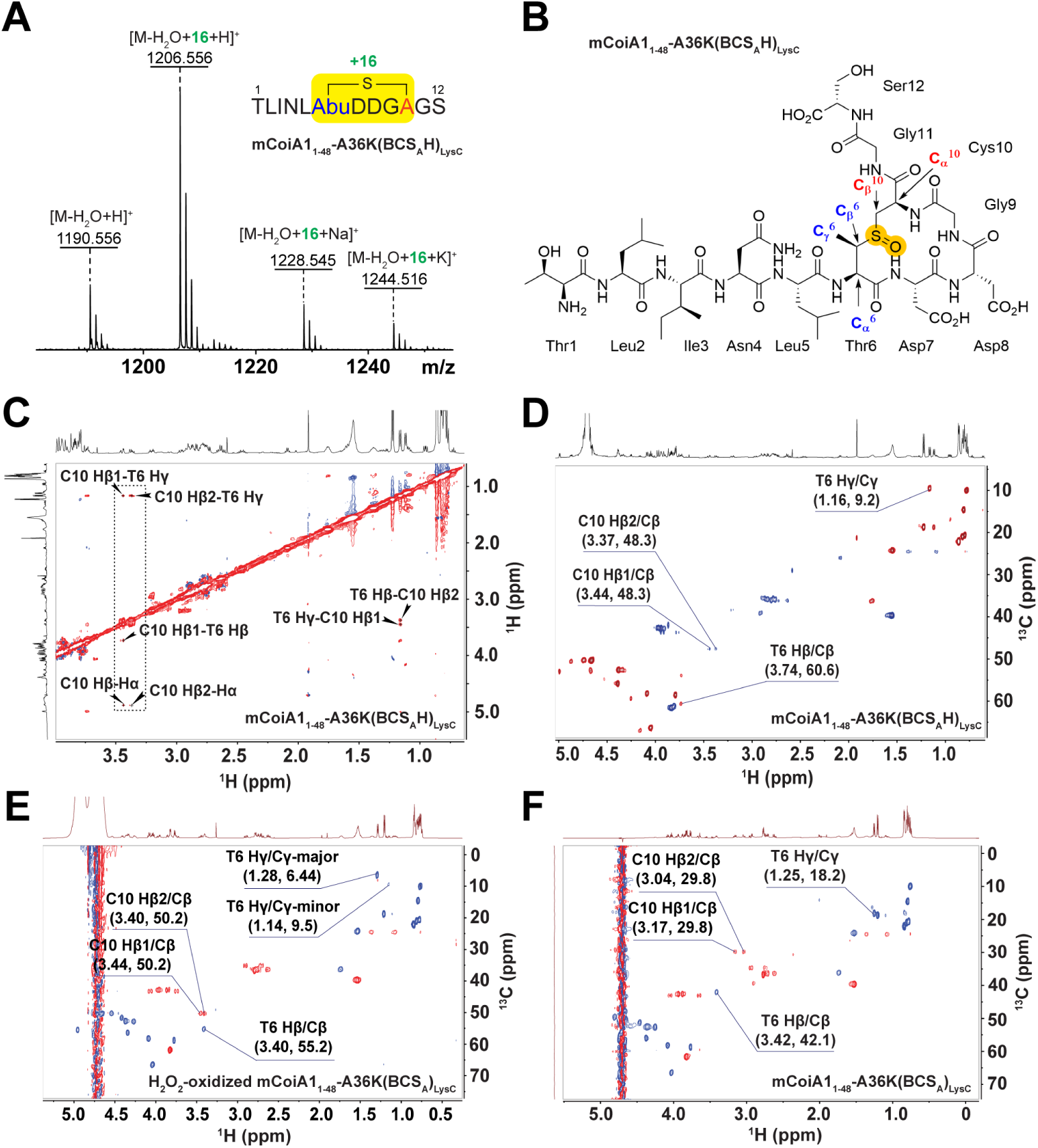
(A) MALDI-TOF MS analysis of LysC-digested His_6_-SUMO-mCoiA1_1-48_-A36K(BCS_A_H). (B) Molecular structure of mCoiA1_1-48_-A36K(BCS_A_H) after LysC digestion. (C) ^1^H-^1^H NOESY spectrum of LysC-digested mCoiA1_1-48_-A36K(BCS_A_H) in 100% D_2_O. Cross-peaks between the β-protons of the former Thr6 and Cys10, as well as between γ-protons of former Thr6 and β-protons of former Cys10 are annotated in the dashed rectangle. (D) ^1^H-^13^C HSQC spectrum of LysC-digested mCoiA1_1-48_-A36K(BCS_A_H) in 100% D_2_O. Cross-peaks with annotation exhibit significant deviations in both ^1^H and ^13^C chemical shifts from the typical values of Thr, Cys, and MeLan residues, indicating their involvement in an oxidized methyllanthionine. (E) ^1^H-^13^C HSQC spectrum of H_2_O_2_-oxidized mCoiA1_1-48_-A36K(BCS_A_)_LysC_ in 90% H_2_O and 10% D_2_O. Two distinct sets of ¹H and ¹³C chemical shifts for the γ-position of former Thr6 were detected. (F) ^1^H-^13^C HSQC spectrum of mCoiA1_1-48_-A36K(BCS_A_)_LysC_ in 90% H_2_O and 10% D_2_O. Complete cross-peak assignments of ^1^H-^13^C HSQC spectra are annotated in Figure S9, and S15-S16, and are summarized in Table S5-S6. For calculated and observed m/z values of molecular ions, see Table S7.

To further confirm the formation of a methyllanthionine sulfoxide and assess if the CoiH transformation is stereoselective, we prepared a large quantity of mCoiA1_1-48_-A36K(BCS_A_)_LysC_ as a control and H_2_O_2_-oxidized mCoiA1_1-48_-A36K(BCS_A_)_LysC_ as a mimic of the CoiH-modified peptide for NMR analysis (Figures S10-S16). For the H_2_O_2_-oxidized peptide, NOESY cross peaks between former Thr6 and former Cys10 were observed, confirming that the methyllanthionine bond remained intact (Figure S14). Upon oxidation by H_2_O_2_, the ^13^C chemical shifts of the β-carbons shifted substantially downfield, from 42.1 ppm to 55.2 ppm for the former Thr6, and from 29.8 ppm to 50.2 ppm for the former Cys10 (Figure 5E-F), which is in agreement with the NMR data of the CoiH-modified product. Notably, the H₂O₂-treated peptide exhibited two distinct sets of ¹H and ¹³C chemical shifts for the γ-position of former Thr6, suggesting the formation of two diastereomers with different stereochemical configurations at the sulfoxide group (Figure 5E and Table S6). Importantly, the Hγ and Cγ chemical shifts of former Thr6 in CoiH-modified peptide closely matched those of the minor diastereomer in the chemically oxidized peptide, which was present at a major-to-minor ratio of 85:15 (Figure S17). These findings indicate the CoiH-catalyzed sulfoxidation is stereoselective and strongly argues against spurious oxidation.

We also evaluated the possibility of L-isoaspartate formation at the second Asp residue in the A-ring of CoiA1, in which the methyltransferase domain of CoiS_A_ may be involved. The ^1^H-^1^H NOESY spectrum of mCoiA1_1-48_-A36K(BCS_A_H)_LysC_ showed a single cross-peak between the amide proton (HN) of Gly9 and the α-proton of Asp8. The absence of a correlation between Gly9 HN and Asp8 Hβ rules out an isoaspartyl linkage, confirming that Asp8 is a standard L-aspartate (Figure S18). Thus, it appears that CoiS_A_ is not active with the truncated CoiA1_1-48_ substrate.

Overall, together with the mass spectrometry results showing a 16 Da mass addition on the N-terminal methyllanthionine, the NMR data obtained from all three peptide samples strongly support the assignment of a sulfoxide group at the thioether linkage site in CoiH-modified CoiA1_1-48_ and confirm that CoiH is an oxygenase catalyzing the site- and stereoselective oxidation of the sulfur of the methyllanthionine A-ring.

### Bioinformatic analysis of CoiH and its homologs

Several enzymes responsible for sulfoxide formation during RiPP biosynthesis have been identified. For example, GarO catalyzes the formation of the sulfoxide moiety in the class II lanthipeptide actagardine,^18,19^ whereas SolS is responsible for sulfoxidation of the labionin residue in deoxylabiomycin.^26^ In fungal RiPP biosynthesis, UstF1 participates in the installation of a sulfoxide group during ustiloxin maturation.^27^ Notably, all of these enzymes belong to the flavin adenine dinucleotide (FAD)-dependent oxygenase family. In contrast, CoiH lacks detectable sequence similarity to these enzymes (Table S8) and a conservation analysis of CoiH and 132 homologs demonstrates that they do not contain any recognizable FAD-binding motifs (Figure S19 and Table S9). Additionally, no significant sequence similarity was detected between CoiH and the non-heme iron-dependent sulfoxide synthases EgtB^28^ and OvoA^29^ (Table S8). CoiH and its homologs also did not share any conserved sequence motifs typically associated with other known oxygenase families, including cytochrome P450s, Rieske oxygenases, or Fe(II)/α-ketoglutarate-dependent oxygenases (Figure S19 and Table S9). BLAST searches with CoiH as query retrieved only uncharacterized homologs of CoiH. An AlphaFold 3 model of the enzyme is shown in Figure S20. Structure-based similarity searches (Foldseek^30^ and DALI^31^) did not detect obvious similarity to proteins with experimentally validated functions. Thus, CoiH and its homologs constitute a separate enzyme family in terms of sequence, structure, and function.

PrankWeb^32^ predicts a putative ligand-binding pocket that contains several highly conserved residues in CoiH (Figure S21 and Table S10). Site-directed mutagenesis and subsequent co-expression studies demonstrated that Asp59, Asn97, Arg100 and Leu332 that line this pocket are essential for activity, consistent with this pocket constituting the enzyme’s active site (Figures S22-S23). Because we have not yet been able to purify CoiH because of solubility and instability issues, we cannot evaluate the expression level and solubility of the CoiH variants. Therefore, we acknowledge that the loss of activity observed for the CoiH variants could potentially result from changes in protein expression, stability, solubility, or folding rather than a direct effect on catalysis. Collectively, these analyses suggest that CoiH does not belong to any currently recognized oxygenase family and may represent a previously uncharacterized enzyme family that catalyzes sulfoxide formation through a distinct mechanism.

### CoiH homologs are associated with lanthivirin BGCs and catalyze sulfoxide formation

BLAST analysis of CoiH identified 3937 homologous proteins, which were subsequently subjected to genome-mining analysis using RODEO.^33^ The vast majority of homologs were encoded within conserved class I lanthipeptide BGC architectures. More than half of these BGCs belong to the *coi*-like BGC family. Other BGCs only differ from *coi*-like BGCs in that either the LanB dehydratase lacks the glutamyl lyase domain and only contains the glutamylation domain or additional tailoring enzymes are encoded (see below). To provide a better overview of the genome mining results, we generated a Sequence Similarity Network (SSN) based on the CoiH homologs using the tools of the Enzyme Function Initiative,^34,35^ which revealed multiple well-separated clusters (Figure 6A). CoiH belongs to cluster 11, whose members as well as the proteins in clusters 1, 2, 3, 4 and 5 are predominantly associated with *coi*-like lanthipeptide BGCs (Figure S24). In contrast, proteins from clusters 6, 7, 9 and 10 are linked to BGCs encoding truncated LanB dehydratases that lack a canonical GL domain (Figure S24). Additionally, the BGCs of cluster 12 encode a TauD/TfdA family dioxygenase which may lead to a new modification of the substrate. The correlation between SSN clustering of CoiH orthologs and BGC architecture suggests that the tailoring enzyme family related to CoiH has co-evolved with its associated biosynthetic pathways.

**Figure 6.**
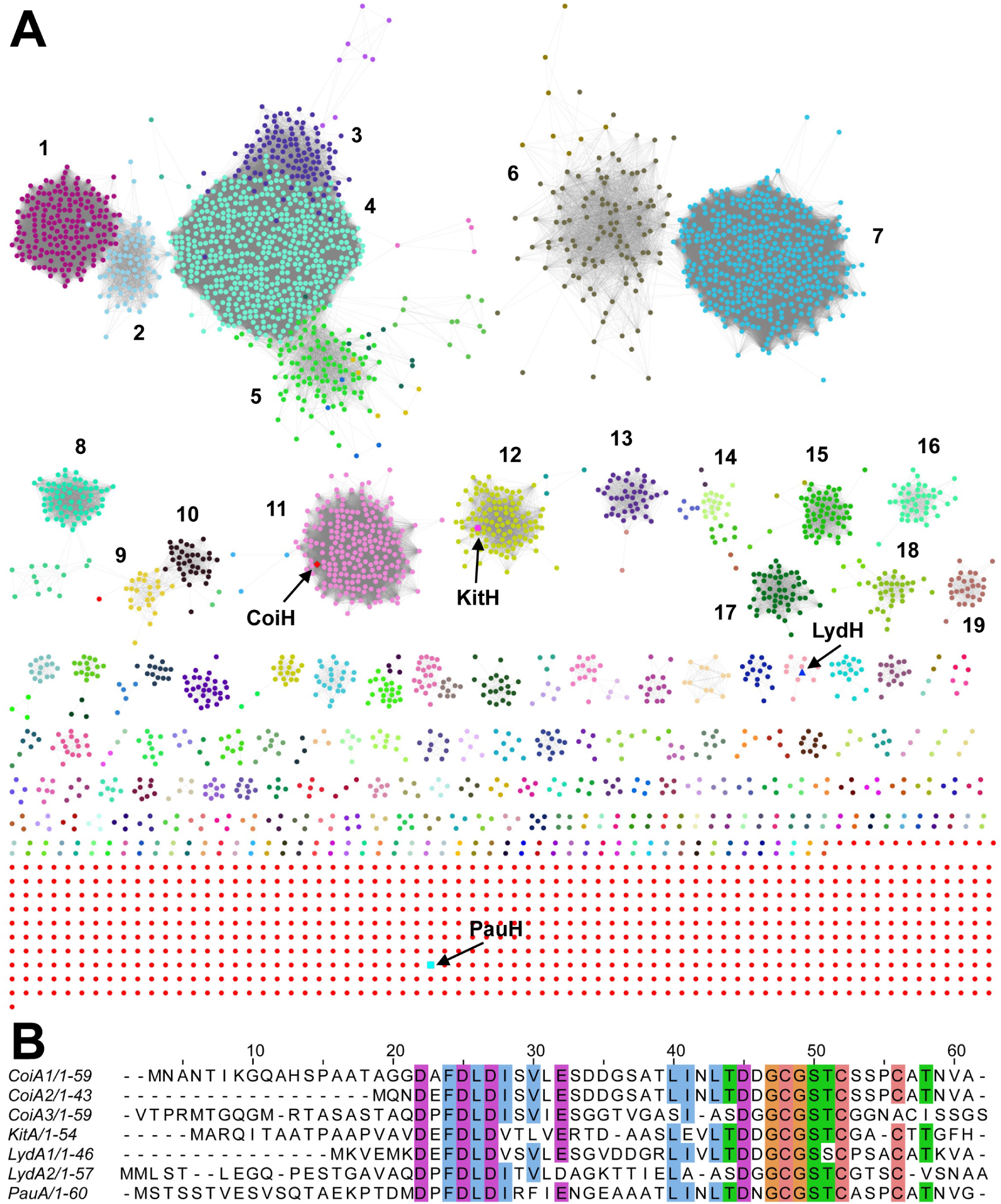
CoiH homologs and selected substrates. (A) SSN analysis of CoiH and its homologs visualized using Cytoscape.^36^ The nineteen largest clusters are annotated 1-19. Proteins investigated in this study (CoiH, KitH, LydH, and PauH) are highlighted using arrows and distinct node shapes. KitH, LydH, and PauH are from *Kitasatospora* sp. NPDC091335, *Streptomyces lydicus*, and *Phytohabitans aurantiacus*, respectively. SSN parameters: alignment score = 140, Protein length = 100-1000 aa, E value = -5. For the architectures of BGCs associated with distinct SSN clusters, see Figure S24. The cytoscape file for the SSN is provided as Supporting Information. (B) Sequence alignment of selected precursor peptides encoded in LanH-associated BGCs. KitA is the substrate of KitH; LydA1 and LydA2 are the substrates of LydH; PauA is the substrate of PauH.

We selected three CoiH homologs KitH (*Kitasatospora* sp. NPDC091335), LydH (*Streptomyces lydicus*), and PauH (*Phytohabitans aurantiacus*) from distinct clusters in the SSN (Figure 6A). These homologs were chosen because of the substantial sequence diversity (42.05 – 54.7% sequence identity to CoiH) and because their substrates all have Thr-Asp-Asp/Asn-Gly-Cys sequences in their substrates that would form the A-rings in their products (Figure 6B). Thus, we anticipated that these enzymes might act on the mCoiA1_1-48_(BCS_A_) peptide. We coexpressed the three enzymes with CoiA1_1-48_ and CoiBCS_A_ in *E. coli.* All three proteins exhibited the same activity and effected methyllanthionine oxidation (Figure S25), albeit not all with the same efficiency. This observation shows that homologs from distinct evolutionary branches retain both catalytic activity and substrate compatibility, which suggests strong evolutionary pressure to maintain the sulfoxidation step across diverse biosynthetic pathways. We attempted to express and purify CoiH, KitH, LydH and PauH for in vitro assays but were not successful due to their poor solubility and instability that precluded isolation.

## Discussion

The studies presented here report on a previously uncharacterized enzyme family. In keeping with the long-standing general nomenclature for lanthipeptides,^37^ we had previously termed this family LanH (e.g. CoiH)^7^ without knowing its enzymatic activity. Here we demonstrate that they catalyze the oxidation of a methyllanthionine to the corresponding sulfoxide. Shomar et al. recently termed these enzymes LvnF (Lvn for lanthivirin) based on the bioactivity of the BGCs (and presumably their products),^10^ but LanF had already been taken in the lanthipeptide nomenclature for transporters.^37^ We feel that LanH and LvnF are both useful terms in different contexts (lanthipeptide biosynthetic enzyme activity and final product bioactivity, respectively).

A related oxidation has been reported previously for the class II lanthipeptide actagardine for which a luciferase-like flavin-dependent enzyme carries out the oxidation of one of the methyllanthionines.^17–19^ The class III lanthipeptides solabiomycins also contain a sulfoxide modification of a labionin crosslink introduced by a flavin dependent enzyme.^26^ For neither of these two peptides is the stereochemistry of the sulfoxide established and our current data also does not resolve the stereochemistry of the CoiH-generated sulfoxide. Another RiPP that contains a sulfoxide is α-aminitin produced by lethal mushrooms. The biosynthetic enzyme has not been definitively established although a flavin enzyme is known to be involved in its biosynthesis.^38^ The sulfoxide group was shown to be essential for antibacterial activity of the solabiomycins,^26^ and the absence of the sulfoxide group in α-aminitin greatly reduces its activity.^39^ We therefore infer that sulfoxide formation is also likely important for the anti-phage activity of the product of the *coi* BGC. In contrast to these previously reported sulfoxide forming enzymes, the LanH enzymes do not appear to bind flavin based on AlphaFold predictions, although we were not able to confirm this hypothesis by in vitro studies because of insolubility. The AlphaFold model and the sequence alignment also do not suggest that these enzymes contain a conserved metal binding site. Thus, we tentatively propose that the LanH proteins may carry out oxidation reactions without the help of a metal or organic cofactor to overcome the kinetic barrier of reactions of triplet oxygen,^40–42^ but their catalytic mechanisms and cofactor requirements remain to be experimentally established. An alternative model that we cannot rule out is that in *E. coli*, small amounts of hydrogen peroxide may be present that are used by the enzyme as the oxidant instead of molecular oxygen. ^18^O_2_-labelling experiments in culture are unlikely to be able to distinguish between molecular oxygen and H_2_O_2_ derived from molecular oxygen as the direct oxidant. Therefore, future in vitro biochemical studies will be necessary to identify the oxygen source for sulfoxidation once the solubility and purification issues of LanH enzymes are resolved.

The antiphage activity of the product of the *coi* BGC has been demonstrated in a previous study,^10^ but the structure of the product was unresolved. With the characterization of the activity of CoiH, the reactions catalyzed by all modification enzymes have been revealed, but some questions remain regarding the final structure of the active peptide. Based on a study on the CoiS_A_ ortholog OlvS_A_ from *Streptomyces olivaceus*,^9^ the methyltransferase domain of CoiS_A_ methylates the side chain of the second Asp in the A-ring of the cyclized peptide. This methylation is followed by aspartimide formation and hydrolytic ring opening to form isoAsp based on NMR analysis of the product. Since the study on OlvS_A_,^9^ other members of this *O*-methyltransferase family have been shown to generate stable aspartimides in a variety of RiPPs.^51–53^ Our NMR experiments showed that isoAsp was not formed in the CoiBCS_A_-modified truncated CoiA1 peptide, but this possibility remains to be investigated in full-length CoiA1. Also still unclear is whether the peptide needs to be proteolytically processed for the anti-phage activity as is seen in most other RiPPs.^2^ The *coi* BGC does not encode a protease, but sometimes the protease(s) involved in RiPP biosynthesis can be encoded remotely.^54^

The lanthipeptides that have been suggested to impart anti-phage activity (lanthivirins) have been divided into 10 clades.^10^ The BGCs of six of these clades (clades 2, 5, 6, and 8-10) contain CoiH homologs, suggesting that the sulfoxide modification is found in about half of the anti-phage lanthipeptides. The *olv* BGC belongs to clade 7 that does not encode a *syn* GL domain, and other lanthivirin BGCs do not contain an *anti*-GL domain (Figure S24). Therefore, the products of the lanthivirin BGCs will contain remarkably diverse stereochemistry of the thioether rings.^7^ These collective considerations illustrate that the lanthivirins display substantial structural diversity. Potentially, the sulfoxide moiety and the different crosslink stereochemistry may be involved in conferring phage specificity. We considered assessing whether the CoiH-modified peptide produced heterologously in *E. coli* exhibits anti-phage activity and whether the absence of the sulfoxide modification affects this activity. However, Shomar et al. proposed that the anti-phage activity of lanthivirin may involve an intracellular mechanism of action.^10^ Given that the boundary between the leader and core peptides has not yet been identified and that the purified lanthivirin produced in *E. coli* is unlikely to enter *Streptomyces* cells, evaluating its anti-phage activity using the heterologously produced peptide is not possible at this stage.

## Conclusion

The *coi* BGC in *S. coelicolor* was shown previously to impart a phage resistance phenotype. The BGC encodes a protein of unknown function CoiH that we show in this work selectively oxidizes the thioether in the A-ring of the lanthipeptide product to the corresponding sulfoxide. Several tested orthologs from homologous BGCs in other Actinobacteria catalyze the same reaction. Structure prediction models suggest that the enzyme does not possess a canonical cofactor or metal binding sequences. Therefore, the catalytic mechanism of CoiH and the identity of the oxidant remain unclear. Further in vitro studies with purified LanH will be necessary to determine whether cofactors or other cellular components are required for stereoselective LanH-catalyzed sulfoxidation.

## Supporting information

Cytoscape file for Fig 6

## ASSOCIATED CONTENT

### Supporting Information

Experimental procedures, peptide characterization data, mass spectrometry data and NMR spectra, AlphaFold 3 modeling, Cytoscape file, and supplementary figures and tables.

## Funding

This manuscript is the result of funding in whole or in part by the National Institutes of Health (NIH; grant R37 GM058822 and R35 GM164113 to W.A.v.d.D.). It is subject to the NIH Public Access Policy. Through acceptance of this federal funding, NIH has been given a right to make this manuscript publicly available in PubMed Central upon the Official Date of Publication, as defined by NIH. A Bruker UltrafleXtreme mass spectrometer used was purchased with support from the Roy J. Carver Charitable Trust (Grant No. 22-5622).

W.A.v.d.D. is an Investigator of the Howard Hughes Medical Institute.

## Notes

The authors declare no competing financial interest.

## Acknowledgements

We thank D. T. Nguyen and C. Padhi for assistance with LC-MS/MS experiments.

## References

(1) Ortega, M. A.; van der Donk, W. A. New insights into the biosynthetic logic of ribosomally synthesized and post-translationally modified peptide natural products. Cell Chem. Biol. 2016, 23, 31–44.

(2) Montalbán-López, M.; Scott, T. A.; Ramesh, S.; Rahman, I. R.; van Heel, A. J.; Viel, J. H.; Bandarian, V.; Dittmann, E.; Genilloud, O.; Goto, Y.; Grande Burgos, M. J.; Hill, C.; Kim, S.; Koehnke, J.; Latham, J. A.; Link, A. J.; Martínez, B.; Nair, S. K.; Nicolet, Y.; Rebuffat, S.; Sahl, H.-G.; Sareen, D.; Schmidt, E. W.; Schmitt, L.; Severinov, K.; Süssmuth, R. D.; Truman, A. W.; Wang, H.; Weng, J.-K.; van Wezel, G. P.; Zhang, Q.; Zhong, J.; Piel, J.; Mitchell, D. A.; Kuipers, O. P.; van der Donk, W. A. New developments in RiPP discovery, enzymology and engineering. Nat. Prod. Rep. 2021, 38, 130–239.

(3) Repka, L. M.; Chekan, J. R.; Nair, S. K.; van der Donk, W. A. Mechanistic understanding of lanthipeptide biosynthetic enzymes. Chem. Rev. 2017, 117, 5457–520.

(4) Ortega, M. A.; Hao, Y.; Zhang, Q.; Walker, M. C.; van der Donk, W. A.; Nair, S. K. Structure and mechanism of the tRNA-dependent lantibiotic dehydratase NisB. Nature 2015, 517, 509–12.

(5) Zhang, Q.; Yu, Y.; Velásquez, J. E.; van der Donk, W. A. Evolution of lanthipeptide synthetases. Proc. Natl. Acad. Sci. U. S. A. 2012, 109, 18361–6.

(6) Sarksian, R.; Hegemann, J. D.; Simon, M. A.; Acedo, J. Z.; van der Donk, W. A. Unexpected methyllanthionine stereochemistry in the morphogenetic lanthipeptide SapT. J. Am. Chem. Soc. 2022, 144, 6373–82.

(7) Sarksian, R.; van der Donk, W. A. Divergent evolution of lanthipeptide stereochemistry. ACS Chem. Biol. 2022, 17, 2551–8.

(8) Sarksian, R.; Zhu, L.; van der Donk, W. A. *syn*-Elimination of glutamylated threonine in lanthipeptide biosynthesis. Chem. Commun. 2023, 59, 1165–8.

(9) Acedo, J. Z.; Bothwell, I. R.; An, L.; Trouth, A.; Frazier, C.; van der Donk, W. A. *O*- methyltransferase-mediated incorporation of a β-amino acid in lanthipeptides. J. Am. Chem. Soc. 2019, 141, 16790–801.

(10) Shomar, H.; Guillaume, M.; Tesson, F.; Conte, V.; Zhang, L.; Du, C.; Ongenae, V.; Loubat, A.; Le Bot, M.; Georjon, H.; Mordret, E.; Rozen, D.; Briegel, A.; Libis, V.; Claessen, D.; Li, Y.; van Wezel, G. P.; Bernheim, A. A family of lanthipeptides with anti-phage function. Cell Host Microbe 2026, 34, 1591–602.

(11) Ökesli, A.; Cooper, L. E.; Fogle, E. J.; van der Donk, W. A. Nine post-translational modifications during the biosynthesis of cinnamycin. J. Am. Chem. Soc. 2011, 133, 13753–60.

(12) Zhang, C.; Seyedsayamdost, M. R. CanE, an iron/2-oxoglutarate-dependent lasso peptide hydroxylase from *Streptomyces canus*. ACS Chem. Biol. 2020, 15, 890–4.

(13) Morishita, Y.; Ma, S.; De La Mora, E.; Li, H.; Chen, H.; Ji, X.; Usclat, A.; Amara, P.; Sugiyama, R.; Tooh, Y. W.; Gunawan, G.; Pérard, J.; Nicolet, Y.; Zhang, Q.; Morinaka, B. I. Fused radical SAM and αKG-HExxH domain proteins contain a distinct structural fold and catalyse cyclophane formation and β-hydroxylation. Nat. Chem. 2024, 16, 1882–93.

(14) Kim, H. W.; Kang, S.; Kim, S.; Lee, H.; Hur, Y.; Song, W. J.; Oh, D. C.; Kim, S. Discovery of a two-step enzyme cascade converting aspartate to aminomalonate in peptide natural product biosynthesis. J. Am. Chem. Soc. 2025, 147, 20909–18.

(15) Ouyang, Y.; Yu, Y.; Zhu, L.; Nguyen, D. T.; van der Donk, W. A. Oxidative peptide backbone cleavage by a HEXXH enzyme during RiPP biosynthesis. J. Am. Chem. Soc. 2026, 148, 3551–61.

(16) Cha, L.; Qian, C.; Padhi, C.; Zhu, L.; van der Donk, W. A. Discovery and biosynthesis of nitrilobacillins by post-translational introduction of C-terminal nitrile Groups. J. Am. Chem. Soc. 2026, 148, 27430–41.

(17) Zimmermann, N.; Metzger, J. W.; Jung, G. The tetracyclic lantibiotic actagardine. 1H-NMR and 13C-NMR assignments and revised primary structure. Eur. J. Biochem. 1995, 228, 786–97.

(18) Boakes, S.; Cortés, J.; Appleyard, A. N.; Rudd, B. A.; Dawson, M. J. Organization of the genes encoding the biosynthesis of actagardine and engineering of a variant generation system. Mol. Microbiol. 2009, 72, 1126–36.

(19) Shi, Y.; Bueno, A.; van der Donk, W. A. Heterologous production of the lantibiotic Ala(0)actagardine in *Escherichia coli*. Chem. Commun. 2012, 48, 10966–8.

(20) Kim, G.; Weiss, S. J.; Levine, R. L. Methionine oxidation and reduction in proteins. Biochim. Biophys. Acta 2014, 1840, 901–5.

(21) Nguyen, D. T.; Zhu, L.; Gray, D. L.; Woods, T. J.; Padhi, C.; Flatt, K. M.; Mitchell, D. A.; van der Donk, W. A. Biosynthesis of macrocyclic peptides with C-terminal β-amino-α-keto acid groups by three different metalloenzymes. ACS Cent. Sci. 2024, 10, 1022–32.

(22) Lohans, C. T.; Vederas, J. C. Structural characterization of thioether-bridged bacteriocins. J. Antibiot. 2014, 67, 23–30.

(23) Hayashi, A.; Goto, Y.; Saito, Y.; Suga, H.; Morimoto, J.; Sando, S. Oxidation-guided and collision-induced linearization assists de novo sequencing of thioether macrocyclic peptides. Chem. Commun. 2024, 60, 9436–9.

(24) Weir, E.; Zhu, L.; van der Donk, W. A. Structure and activity of class II lanthipeptides from a thermophilic bacterium. ChemBioChem 2026, 27, e70440.

(25) Dračínský, M.; Pohl, R.; Slavětínská, L.; Buděšínský, M. Observed and calculated 1H and 13C chemical shifts induced by the in situ oxidation of model sulfides to sulfoxides and sulfones. Magn. Reson. Chem. 2010, 48, 718–26.

(26) Ijichi, S.; Hoshino, S.; Asamizu, S.; Onaka, H. SolS-catalyzed sulfoxidation of labionin to solabionin drives antibacterial activity of solabiomycins. Bioorg. Med. Chem. Lett. 2023, 89, 129323.

(27) Ye, Y.; Minami, A.; Igarashi, Y.; Izumikawa, M.; Umemura, M.; Nagano, N.; Machida, M.; Kawahara, T.; Shin-Ya, K.; Gomi, K.; Oikawa, H. Unveiling the biosynthetic pathway of the ribosomally synthesized and post-translationally modified peptide ustiloxin B in filamentous fungi. Angew. Chem. Int. Ed. 2016, 55, 8072-5.

(28) Seebeck, F. P. In vitro reconstitution of Mycobacterial ergothioneine biosynthesis. J Am Chem Soc 2010, 132, 6632–3.

(29) Braunshausen, A.; Seebeck, F. P. Identification and characterization of the first ovothiol biosynthetic enzyme. J Am Chem Soc 2011, 133, 1757–9.

(30) van Kempen, M.; Kim, S. S.; Tumescheit, C.; Mirdita, M.; Lee, J.; Gilchrist, C. L. M.; Söding, J.; Steinegger, M. Fast and accurate protein structure search with Foldseek. Nat. Biotechnol. 2024, 42, 243–6.

(31) Holm, L.; Laiho, A.; Törönen, P.; Salgado, M. DALI shines a light on remote homologs: One hundred discoveries. Protein Sci. 2023, 32, e4519.

(32) Polák, L.; Škoda, P.; Riedlová, K.; Krivák, R.; Novotný, M.; Hoksza, D. PrankWeb 4: a modular web server for protein-ligand binding site prediction and downstream analysis. Nucleic Acids Res. 2025, 53, W466–w71.

(33) Tietz, J. I.; Schwalen, C. J.; Patel, P. S.; Maxson, T.; Blair, P. M.; Tai, H. C.; Zakai, U. I.; Mitchell, D. A. A new genome-mining tool redefines the lasso peptide biosynthetic landscape. Nat. Chem. Biol. 2017, 13, 470–8.

(34) Gerlt, J. A.; Bouvier, J. T.; Davidson, D. B.; Imker, H. J.; Sadkhin, B.; Slater, D. R.; Whalen, K. L. Enzyme Function Initiative-Enzyme Similarity Tool (EFI-EST): A web tool for generating protein sequence similarity networks. Biochim. Biophys. Acta 2015, 1854, 1019–37.

(35) Oberg, N.; Zallot, R.; Gerlt, J. A. EFI-EST, EFI-GNT, and EFI-CGFP: Enzyme Function Initiative (EFI) web resource for genomic enzymology tools. J. Mol. Biol. 2023, 435, 168018.

(36) Shannon, P.; Markiel, A.; Ozier, O.; Baliga, N. S.; Wang, J. T.; Ramage, D.; Amin, N.; Schwikowski, B.; Ideker, T. Cytoscape: a software environment for integrated models of biomolecular interaction networks. Genome Res. 2003, 13, 2498–504.

(37) de Vos, W. M.; Jung, G.; Sahl, H.-G. In Nisin and Novel Lantibiotics; Jung, G., Sahl, H.-G., Eds.; ESCOM: Leiden, 1991, p 457–64.

(38) Luo, H.; Hallen-Adams, H. E.; Lüli, Y.; Sgambelluri, R. M.; Li, X.; Smith, M.; Yang, Z. L.; Martin, F. M. Genes and evolutionary fates of the amanitin biosynthesis pathway in poisonous mushrooms. Proc. Natl Acad. Sci. U. S. A. 2022, 119, e2201113119.

(39) Walton, J. D. *The Cyclic Peptide Toxins of Amanita and Other Poisonous Mushrooms*; Springer International Publishing AG: Cham, Switzerland, 2018.

(40) Thierbach, S.; Bui, N.; Zapp, J.; Chhabra, S. R.; Kappl, R.; Fetzner, S. Substrate-assisted O_2_ activation in a cofactor-independent dioxygenase. Chem. Biol. 2014, 21, 217–25.

(41) Bugg, T. D. How to break the rules of dioxygen activation. Chem. Biol. 2014, 21, 168–9.

(42) Fetzner, S.; Steiner, R. A. Cofactor-independent oxidases and oxygenases. Appl. Microbiol. Biotechnol. 2010, 86, 791–804.

(43) Machovina, M. M.; Usselman, R. J.; DuBois, J. L. Monooxygenase substrates mimic flavin to catalyze cofactorless oxygenations. J. Biol. Chem. 2016, 291, 17816–28.

(44) Hoffarth, E. R.; Caddell Haatveit, K.; Kuatsjah, E.; MacNeil, G. A.; Saroya, S.; Walsby, C. J.; Eltis, L. D.; Houk, K. N.; Garcia-Borràs, M.; Ryan, K. S. A shared mechanistic pathway for pyridoxal phosphate-dependent arginine oxidases. Proc. Natl. Acad. Sci. U. S. A. 2021, 118, e2012591118.

(45) Tseng, C. C.; Vaillancourt, F. H.; Bruner, S. D.; Walsh, C. T. DpgC is a metal- and cofactor-free 3,5-dihydroxyphenylacetyl-CoA 1,2-dioxygenase in the vancomycin biosynthetic pathway. Chem. Biol. 2004, 11, 1195–203.

(46) Bui, S.; Gil-Guerrero, S.; van der Linden, P.; Carpentier, P.; Ceccarelli, M.; Jambrina, P. G.; Steiner, R. A. Evolutionary adaptation from hydrolytic to oxygenolytic catalysis at the α/β-hydrolase fold. Chem. Sci. 2023, 14, 10547–60.

(47) Li, K.; Fielding, E. N.; Condurso, H. L.; Bruner, S. D. Probing the structural basis of oxygen binding in a cofactor-independent dioxygenase. Acta Crystallogr. D Struct. Biol. 2017, 73, 573–80.

(48) Widboom, P. F.; Fielding, E. N.; Liu, Y.; Bruner, S. D. Structural basis for cofactor-independent dioxygenation in vancomycin biosynthesis. Nature 2007, 447, 342–5.

(49) Luther, G. W.; Findlay, A. J.; MacDonald, D. J.; Owings, S. M.; Hanson, T. E.; Beinart, R. A.; Girguis, P. R. Thermodynamics and kinetics of sulfide oxidation by oxygen: a look at inorganically controlled reactions and biologically mediated processes in the environment. Front. Microbiol. 2011, 2, No. 61.

(50) Miller, B. L.; Williams, T. D.; Schöneich, C. Mechanism of sulfoxide formation through reaction of sulfur radical cation complexes with superoxide or hydroxide ion in oxygenated aqueous solution. J. Am. Chem. Soc. 1996, 118, 11014-25.

(51) Cao, L.; Beiser, M.; Koos, J. D.; Orlova, M.; Elashal, H. E.; Schroder, H. V.; Link, A. J. Cellulonodin-2 and lihuanodin: lasso peptides with an aspartimide post-translational modification. J. Am. Chem. Soc. 2021, 143, 11690–702.

(52) Elashal, H. E.; Koos, J. D.; Cheung-Lee, W. L.; Choi, B.; Cao, L.; Richardson, M. A.; White, H. L.; Link, A. J. Biosynthesis and characterization of fuscimiditide, an aspartimidylated graspetide. Nat. Chem. 2022, 14, 1325–34.

(53) Cao, L.; Elashal, H. E.; Link, A. J. Kinetics of aspartimide formation and hydrolysis in lasso peptide lihuanodin. Biochemistry 2023, 62, 695–9.

(54) Eslami, S. M.; van der Donk, W. A. Proteases involved in leader peptide removal during RiPP biosynthesis. ACS Bio. Med. Chem. Au 2024, 4, 20–36.

