## Supporting Information for "A Distinct Oxygenase Family Catalyzing Methyllanthionine Sulfoxide Formation in Anti-phage Class I Lanthipeptides"

All data have been deposited at:

Chen, Jingqi; Zhu, Lingyang; van der Donk, Wilfred (2026), “Data associated with "A New Enzyme Family Catalyzing Methyllanthionine Sulfoxide Formation in Anti-phage Lanthipeptides"”, Mendeley Data, V1, doi: 10.17632/m3xjjfn9cb.1

### Experimental Methods

#### General materials and methods

Unless otherwise specified, all reagents, buffer components, and LB medium were obtained from Fisher Scientific (Hampton, NH). Oligonucleotide primers were synthesized by Integrated DNA Technologies (Coralville, IA), whereas DNA polymerase and other molecular biology enzymes were purchased from New England Biolabs (Ipswich, MA). Plasmid DNA purification and agarose gel DNA recovery were carried out using spin-column kits from Syd Labs (MA) following the manufacturer's instructions. Sequencing of plasmid constructs was performed by Plasmidsaurus (Louisville, KY). Antibiotics and IPTG used for bacterial cultivation and protein expression were obtained from Gold Biotechnology. For routine cloning and plasmid amplification, *E. coli* DH5α was employed, while recombinant proteins and peptides were expressed in *E. coli* BL21(DE3). All plasmids used and created are provided in Table S13. All primers used are shown in Table S14.

#### Plasmid construction

DNA inserts and linearized vectors were generated by PCR, followed by agarose gel electrophoresis and purification using spin-column kits. Gibson assembly reactions were carried out by combining inserts and vectors at a 2:1 molar ratio (insert: vector) with Gibson Assembly Master Mix according to the manufacturer's instructions. The assembled plasmids were used to transform chemically competent *Escherichia coli* DH5α cells and plated on LB agar containing the appropriate antibiotic: 50 μg/mL kanamycin for pRSFDuet-derived constructs, 100 μg/mL ampicillin for pETDuet-derived constructs, or 100 μg/mL streptomycin for pCDFDuet-derived constructs. Individual colonies were cultured in LB medium supplemented with the corresponding antibiotic, and plasmids were purified for DNA sequencing to verify the correct constructs. The $coiS_A$ gene was inserted into the MCS2 site of pETDuet by Gibson Assembly to generate pETDuet-$CoiS_A$. Codon-optimized *kitH*, *lydH*, and *pauH* genes were synthesized by Twist Bioscience (Table S15) and individually cloned into pETDuet-CoiC-$CoiS_A$. Site-directed mutations were generated by PCR-based mutagenesis, and all mutations were confirmed by DNA sequencing.

#### Heterologous expression and purification of CoiA1 and modified CoiA1

*E. coli* BL21 (DE3) cells were co-transformed with expression plasmids in the appropriate combinations to test the in cellulo activities of specific enzymes (see Table S13). Transformants were selected on LB agar plates containing the corresponding antibiotics. Individual colonies were cultured overnight in 50 mL of LB medium supplemented with the appropriate antibiotics and subsequently used to inoculate 1.0 L of LB cultures. Cells were grown at 37 °C with shaking (200 rpm) until an $OD_{600}$ of approximately 0.6 was reached.

Protein expression was induced with 0.8 mM IPTG after the cultures had been chilled on ice for 30 min, followed by incubation at 18 °C with shaking (220 rpm) for 16–20 h. Cells were collected by centrifugation (10,500 x g, 15 min), and the resulting pellets were used for peptide purification.

Cell pellets obtained from 1.0-L cultures were resuspended in 30 mL of lysis buffer (20 mM Tris-HCl, 300 mM NaCl, 10 mM imidazole, 6 M guanidine hydrochloride, pH 8.0) and disrupted by sonication. Insoluble material was removed by centrifugation (18,000 × *g*, 60 min, 4 °C), after which the clarified lysate was loaded onto a 2-mL Ni-NTA agarose column. Following washing with 20 mL of wash buffer (20 mM Tris-HCl, 300 mM NaCl, 30 mM imidazole, 6 M guanidine hydrochloride, pH 8.0), bound peptides were eluted using 6 mL of elution buffer (20 mM Tris-HCl, 300 mM NaCl, 500 mM imidazole, pH 8.0). The eluted peptide samples were transferred into the desired buffer by buffer exchange and subsequently concentrated with Amicon Ultra centrifugal filters (3.0 kDa MWCO). Concentrated peptide samples were subjected to either TEV protease, GluC, AspN, or LysC digestion prior to further treatment. For TEV-mediated His6-SUMO removal, peptide samples were incubated with His-tagged TEV protease in 20 mM Tris-HCl (pH 8.0) at room temperature for 12 h. Following digestion, the reaction mixture was passed through Ni-NTA resin to remove both the His-tag and TEV protease. The released peptides were analyzed by MALDI-TOF MS, and correctly processed products were stored at −20 °C until further use. For GluC or LysC digestion, peptide samples were incubated with enzyme in 50 mM Tris-HCl (pH 8.0) at 37 °C for 2 to18 h.

Before HPLC purification, the digestion mixture was clarified by centrifugation (16,000 x g, 10 min) to remove insoluble material. Peptides were purified on a Jupiter 5u C18 column (5 µm x 2.00 mm x 150 mm) using an Agilent 1260 Infinity III system. Mobile phase A consisted of water containing 0.1% TFA, whereas mobile phase B contained acetonitrile supplemented with 0.1% TFA. Peptides were separated using the following gradient: 0–5 min 5% B, 5–20 min 5–60% B, 20–28 min 60–100% B, 28–30 min 100% B, 30–32 min 5% B, at a flow rate of 1.2 mL $min^{-1}$ with UV detection at 220 nm. Fractions containing the desired peptide were identified by MALDI-TOF MS, pooled, lyophilized, and used for subsequent biochemical assays or NMR analysis.

The mCoiA1$_{1\text{-}48}$-A36K(BCS$_A$) and mCoiA1$_{1\text{-}48}$-A36K(BCS$_A$H) peptides were digested by LysC followed by HPLC purification. The purified mCoiA1$_{1\text{-}48}$-A36K(BCS$_A$H)$_{LysC}$ peptide consisted of two modified forms of the peptide, M-$H_2O$ (~20%) and M-$H_2O$+O (~80%), and this mixture was used directly for NMR experiments. A small amount of non-dehydrated peptide was also detected by NMR.

The mCoiA1$_{1\text{-}48}$-A36K(BCS$_A$)$_{LysC}$ peptide was prepared at a concentration of 0.7 mM and subjected to NMR analysis prior to $H_2O_2$ oxidation. Following $H_2O_2$ treatment, the oxidized mCoiA1$_{1\text{-}48}$-A36K(BCS$_A$)$_{LysC}$ peptide was obtained at a concentration of 0.6 mM and subsequently analyzed by NMR.

**Oxidation of mCoiA1$_{1\text{-}48}$ (BCS$_A$H)$_{TEV}$ and mCoiA1$_{1\text{-}48}$-A36K(BCS$_A$)$_{LysC}$ with $H_2O_2$**

Hydrogen peroxide was added to peptide solutions to give final concentrations of 1 M $H_2O_2$ and 20 µM peptide. Briefly, 20 µL of 15.3% (w/w) $H_2O_2$ was combined with 80 µL of peptide solution (25 µM in 0.25% DMSO/water), and the reaction was allowed to proceed at 37 °C for 4 h. The reaction was subsequently quenched by the addition of sodium pyruvate, which reacts with $H_2O_2$ to form alanine.[1]

**MALDI-TOF MS**

MALDI-TOF mass spectrometric analyses were performed on either a Bruker UltrafleXtreme MALDI-TOF/TOF or a Bruker Autoflex Speed LRF MALDI-TOF instrument operated in positive reflector mode. Prior to analysis, peptide samples were either purified by HPLC or desalted using ZipTip C18 pipette tips (Millipore). For sample preparation, 1 µL of the sample solution was first applied to the MALDI target and allowed to air dry, after which 1 µL of a 50 mg/mL Super-DHB (2,5-dihydroxybenzoic acid and 2-hydroxy-5-methoxybenzoic acid) solution in acetonitrile was spotted onto the same position. Mass spectra were processed and analyzed using Bruker flexAnalysis software.

**LC-MS/MS**

LC–MS/MS analyses were performed using an Agilent 1290 Infinity II LC system coupled to an Agilent 6545B Q-TOF ESI mass spectrometer. Desalted or HPLC-purified peptide samples were separated on a Phenomenex Aeris C18-XB column (150 × 4.6 mm, 100 Å pore size, 2.6 µm particle size) at a flow rate of 0.6 mL $min^{-1}$ using the following gradient: 5% MeCN containing 0.1% formic acid for 3 min, 5–70% over 12 min, and 70–90% over 3 min (aqueous mobile phase contained 0.1% formic acid). MS/MS spectra were acquired in positive-ion mode with a collision energy of 25–35 mV. Fragment ions were first annotated using the Interactive Peptide Spectral Annotator (IPSA)[2] and subsequently verified by manual inspection.

**NMR spectroscopy**

For NMR analysis, the lyophilized peptide was dissolved in either 280 µL of 90% $H_2O$/10% $D_2O$ or 100% $D_2O$ and transferred to a $D_2O$-matched Shigemi tube. One-dimensional $^1$H NMR spectra were acquired on a 750 MHz Agilent VNMRS spectrometer equipped with a 5 mm HCN probe. Two-dimensional $^1$H-$^1$H NOESY, $^1$H-$^1$H TOCSY, and $^1$H-$^{13}$C HSQC spectra were subsequently recorded on a 600 MHz Bruker NEO spectrometer fitted with a 5 mm broadband Prodigy CryoProbe.

**Bioinformatics**

Protein homologs were identified by BLAST searches, the associated biosynthetic gene clusters were analyzed using RODEO2,[3] and structural homology searches were performed using Foldseek[4] and DALI.[5] EFI-EST was used to perform analysis of CoiH and its homologs (full size = 3938 members) with the following parameters: alignment score = 140, Protein length = 100-1000 aa, E value = -5, to generate a sequence similarity network (SSN).[6] Multiple sequence alignment of CoiH homologous proteins (N=133) was generated using MAFFT and used for subsequent sequence analyses.[7] Potential ligand-binding pockets of the AlphaFold-predicted protein structure were identified using PrankWeb with default parameters.[8]

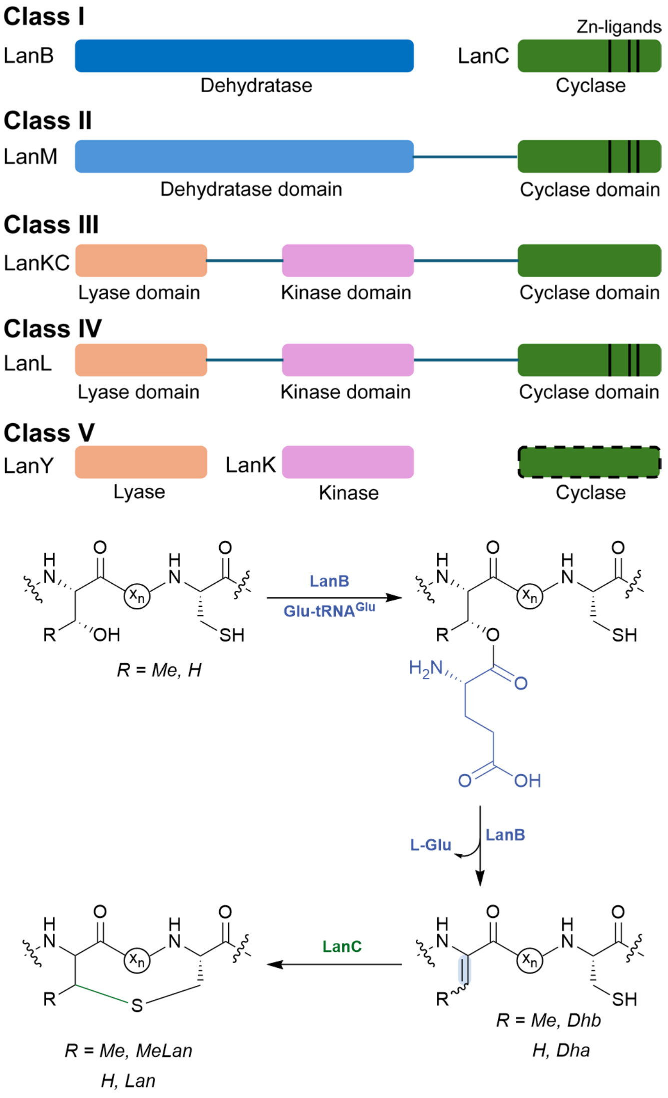

**Figure S1.** Classification of lanthipeptides. They are divided into five classes (I–V) primarily by the biosynthetic enzymes that catalyze dehydration of Ser/Thr residues and subsequent cyclization with Cys residues. The mechanism of biosynthesis of class I lanthipeptides is shown.

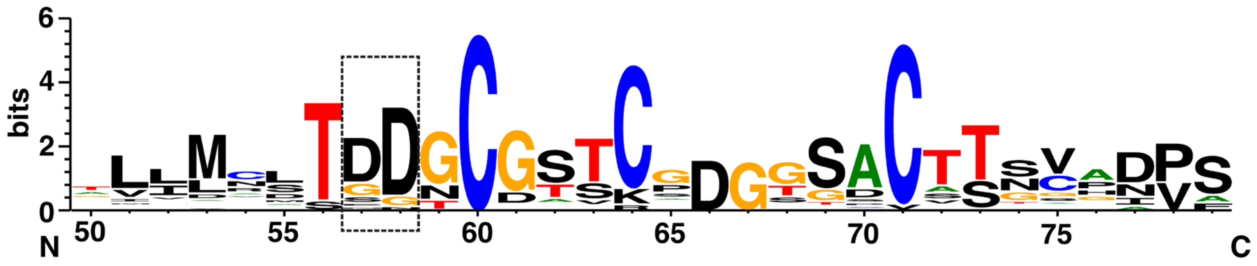

**Figure S2.** Sequence LOGO of CoiA1 and its homologs. The conserved Asp-Asp motif in the thioether A-ring area is highlighted in the dashed rectangle. The LanA sequences were selected from *coi*-like BGCs identified by genome mining (total number of sequences = 316).

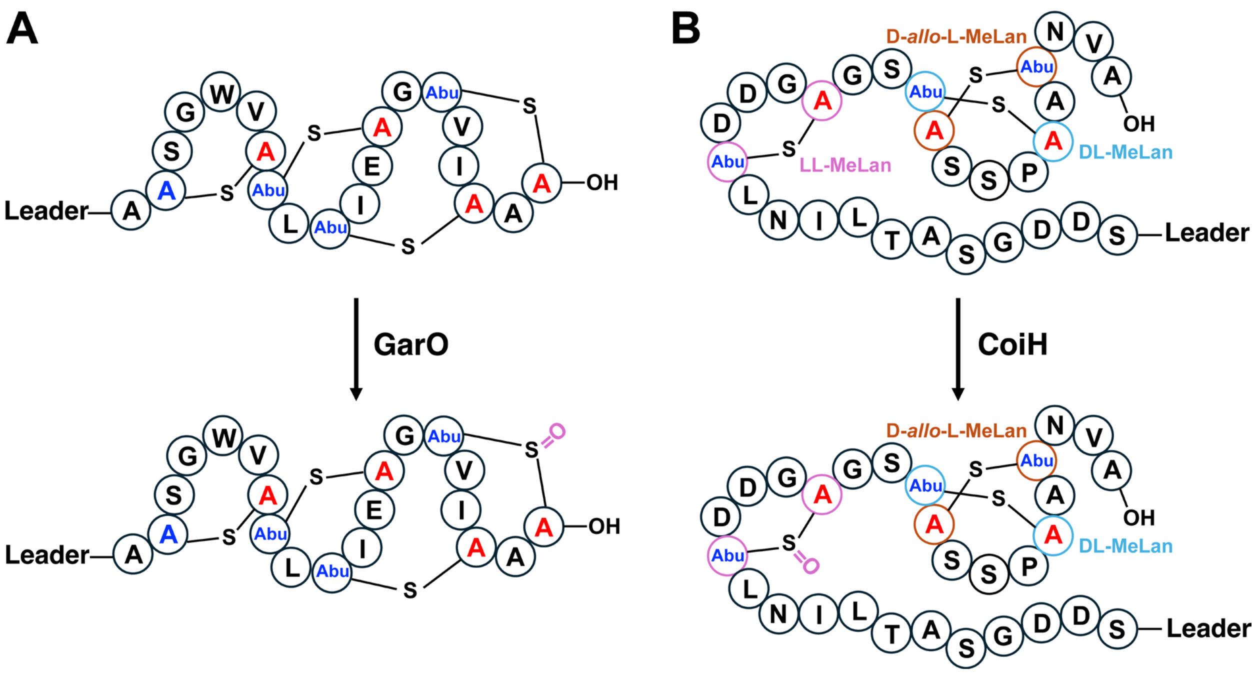

**Figure S3.** The formation of methyllanthionine sulfoxide catalyzed by GarO (experimentally verified) and CoiH (proposed) in lanthipeptide biosynthesis. (A) GarO is a flavin-dependent monooxygenase in actagardine biosynthesis. It catalyzes the selective oxidation of the methyllanthionine sulfur to form methyllanthionine sulfoxide. (B) CoiH is proposed to catalyze a reaction similar to that of GarO, oxidizing the methyllanthionine sulfur to form methyllanthionine sulfoxide in the A-ring.

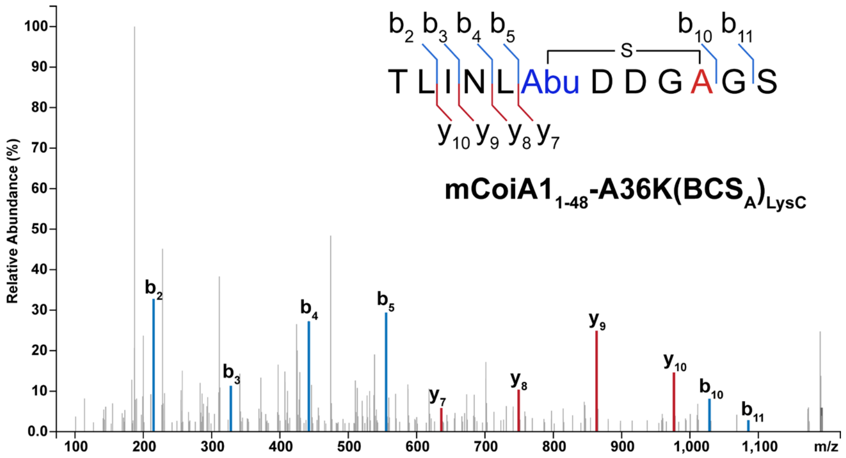

**Figure S4.** LC-MS/MS analysis of mCoiA1$_{1-48}$-A36K(BCS$_A$)$_{LysC}$. The fragmentation pattern is consistent with one MeLan in the CoiBCS$_A$-modified CoiA1$_{1-48}$-A36K peptide. For calculated and observed m/z values of molecular ions, see Table S4.

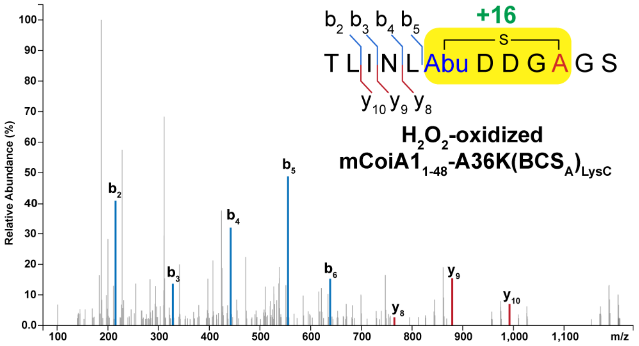

**Figure S5.** LC-MS/MS analysis of the $H_2O_2$-oxidized mCoiA1$_{1-48}$-A36K(BCS$_A$)$_{LysC}$. $H_2O_2$ treatment at 37 °C for 4 h resulted in one +16 Da adduct in the MeLan (yellow-highlighted region). For calculated and observed m/z values of molecular ions, see Table S4.

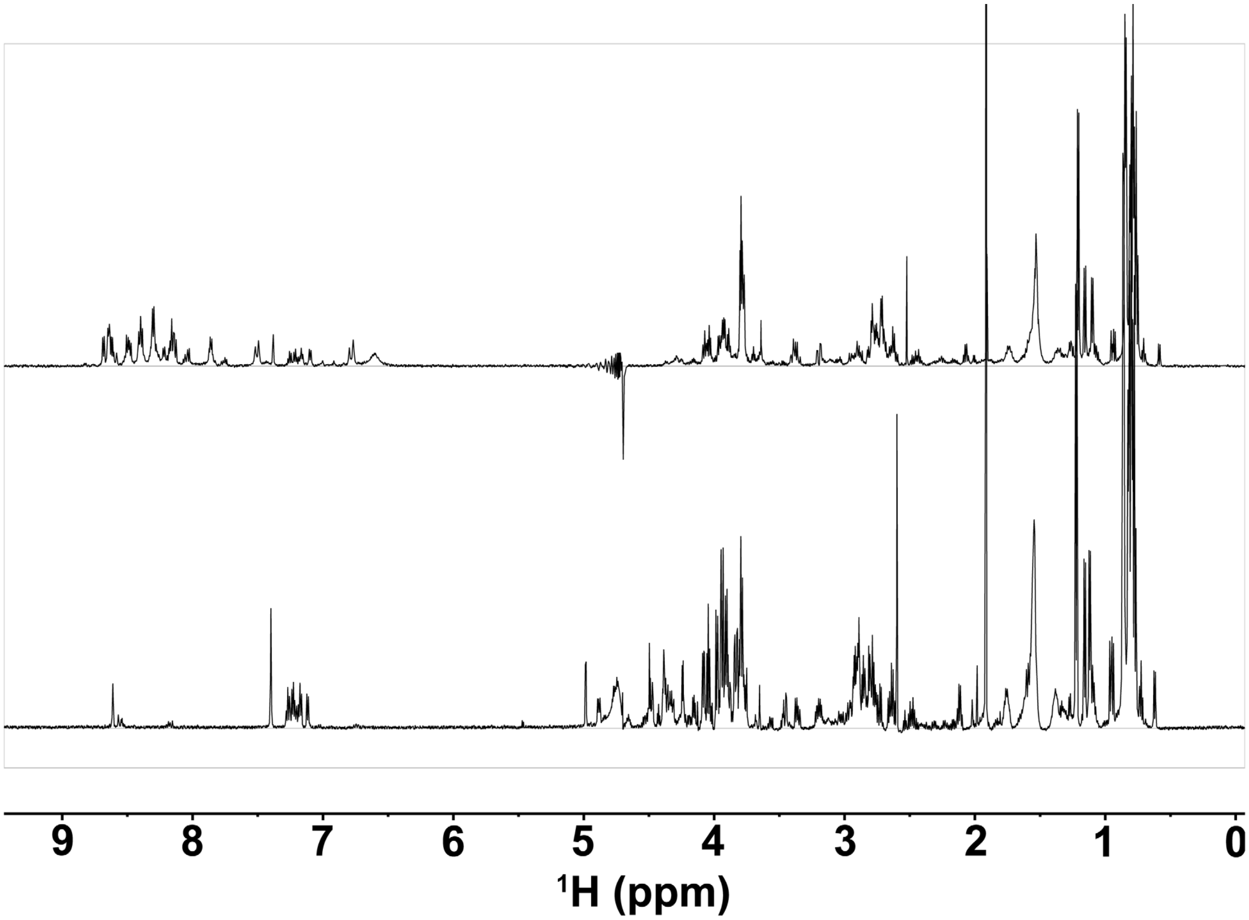

**Figure S6.** $^{1}$H NMR spectra of LysC-digested mCoiA1$_{1\text{-}48}$-A36K(BCS$_A$H). Top, in 90% $H_2O$ and 10% $D_2O$; bottom, in 100% $D_2O$.

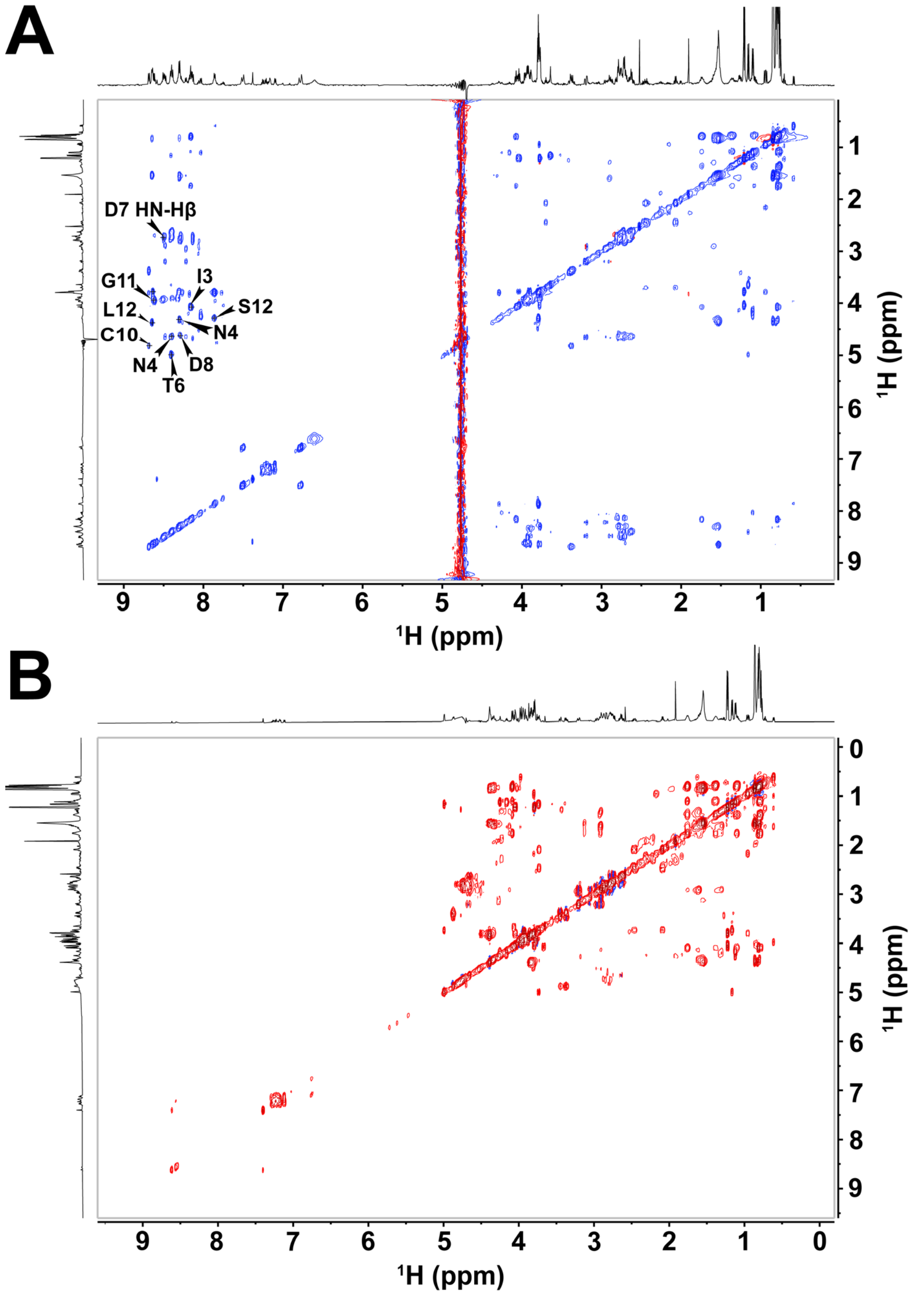

**Figure S7.** (A) $^1H$-$^1H$ TOCSY spectrum of LysC-digested mCoiA1$_{1\text{-}48}$-A36K(BCS$_A$H) in 90% $H_2O$ and 10% $D_2O$. Select amide proton assignments are annotated in the figure. (B) $^1H$-$^1H$ TOCSY spectrum of LysC-digested mCoiA1$_{1\text{-}48}$-A36K(BCS$_A$H) in 100% $D_2O$. Assignments are provided in Table S5.

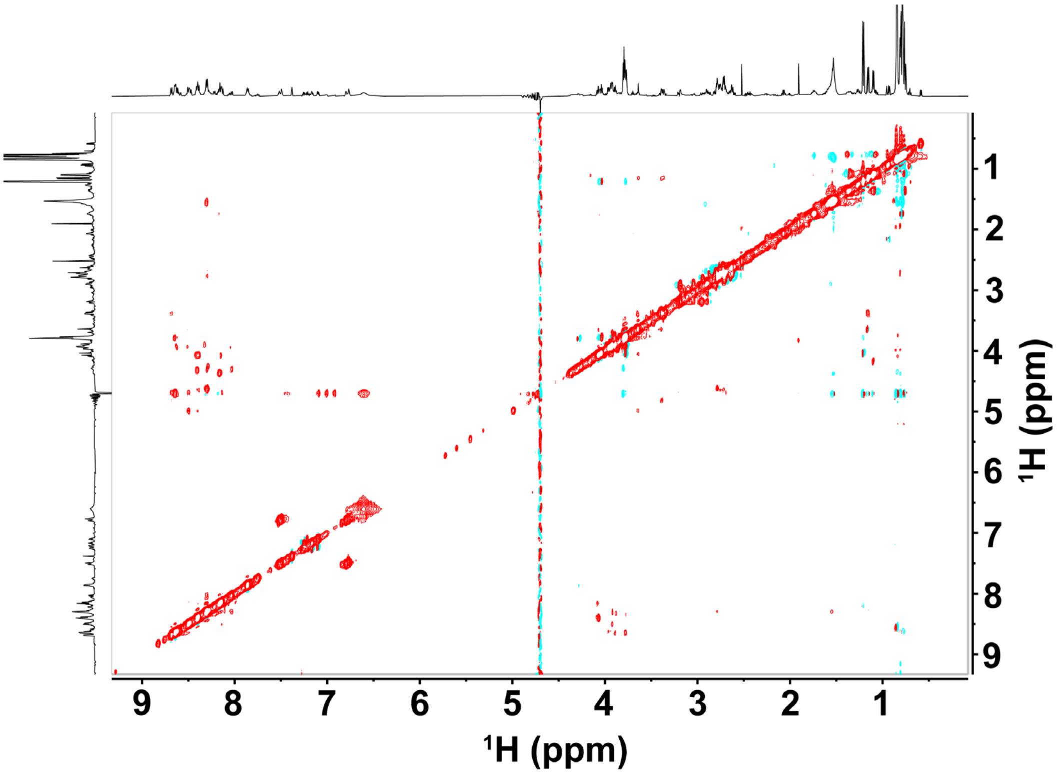

**Figure S8.** $^1$H-$^1$H NOESY spectrum of LysC-digested mCoiA1$_{1\text{-}48}$-A36K(BCS$_A$H) in 90% $H_2O$ and 10% $D_2O$.

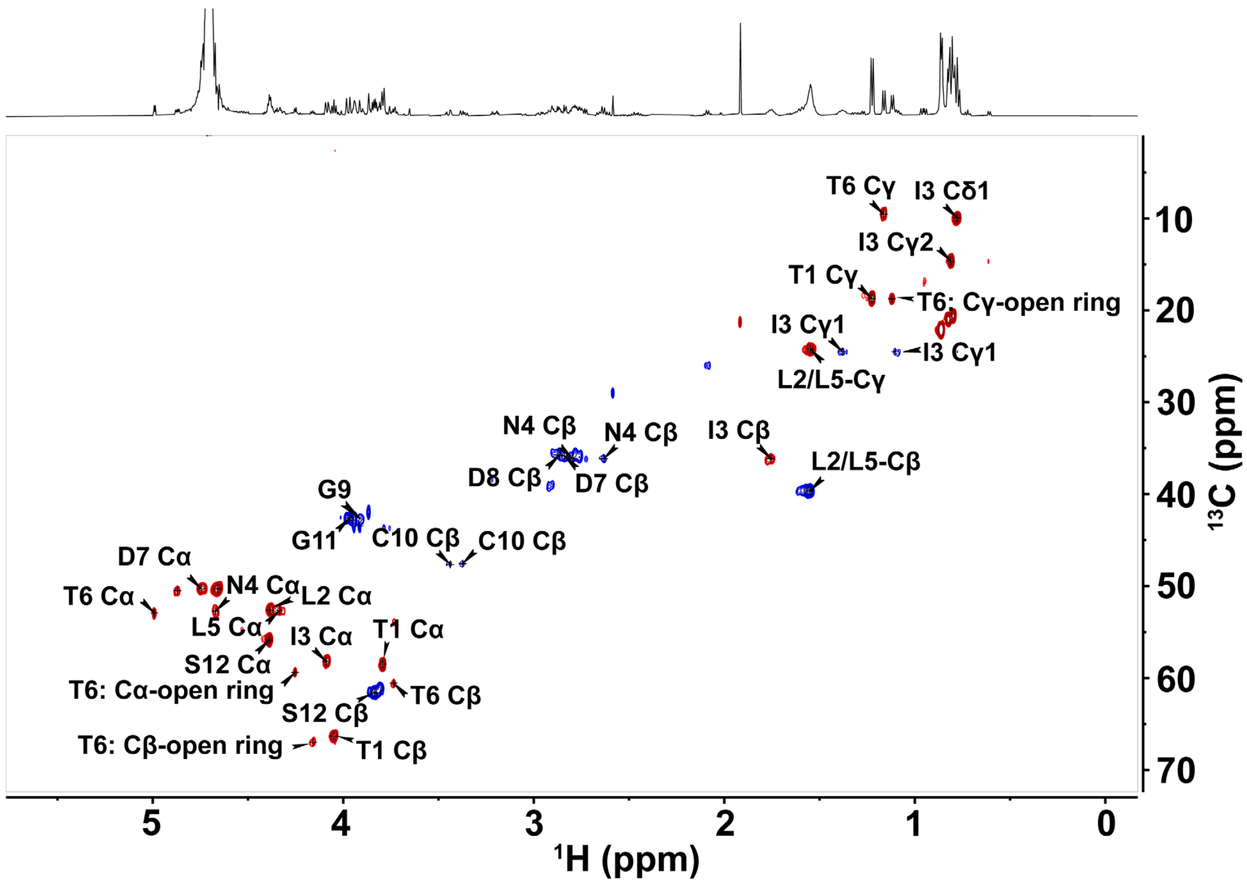

**Figure S9.** $^{1}$H-$^{13}$C HSQC spectrum of LysC-digested mCoiA1$_{1\text{-}48}$-A36K(BCS$_A$H) in 100% $D_2O$. The cross-peaks are annotated in the figure, and their assignments are summarized in Table S5. T6 annotated as open ring peptide is non-dehydrated Thr.

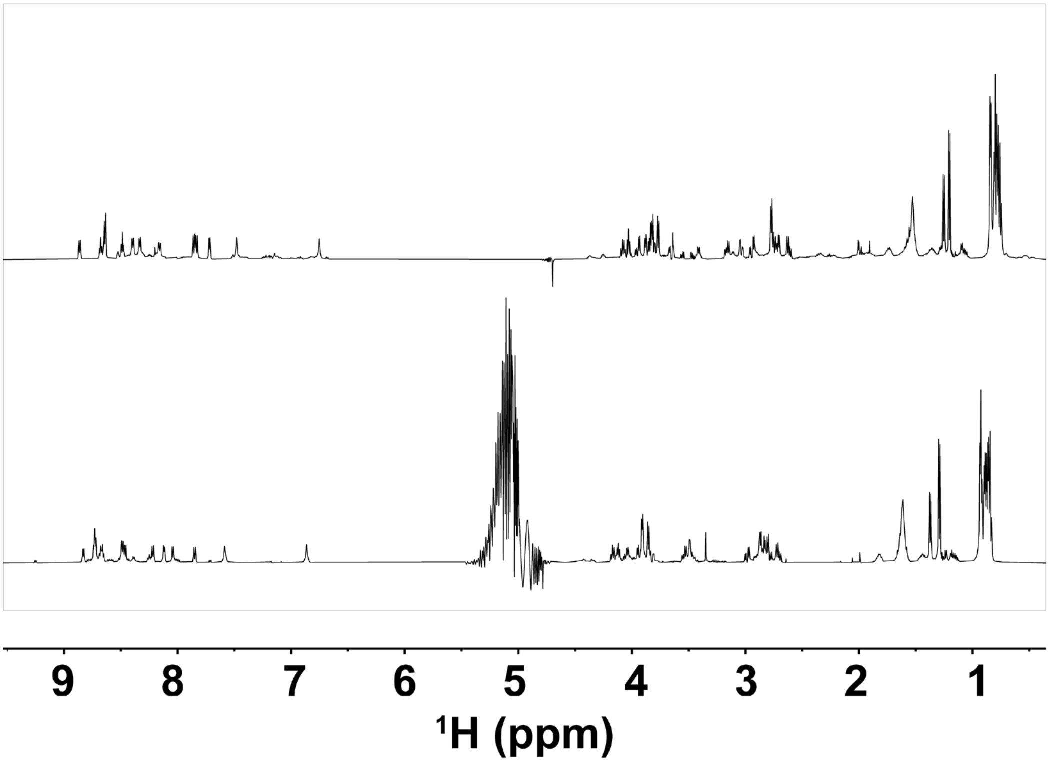

**Figure S10.** ¹H NMR spectra of mCoiA1$_{1\text{-}48}$-A36K(BCS$_{A)}$$_{LysC}$ (top) and $H_2O_2$-oxidized mCoiA1$_{1\text{-}48}$-A36K (BCS$_{A)}$$_{LysC}$ (bottom) in 90% $H_2O$ and 10% $D_2O$.

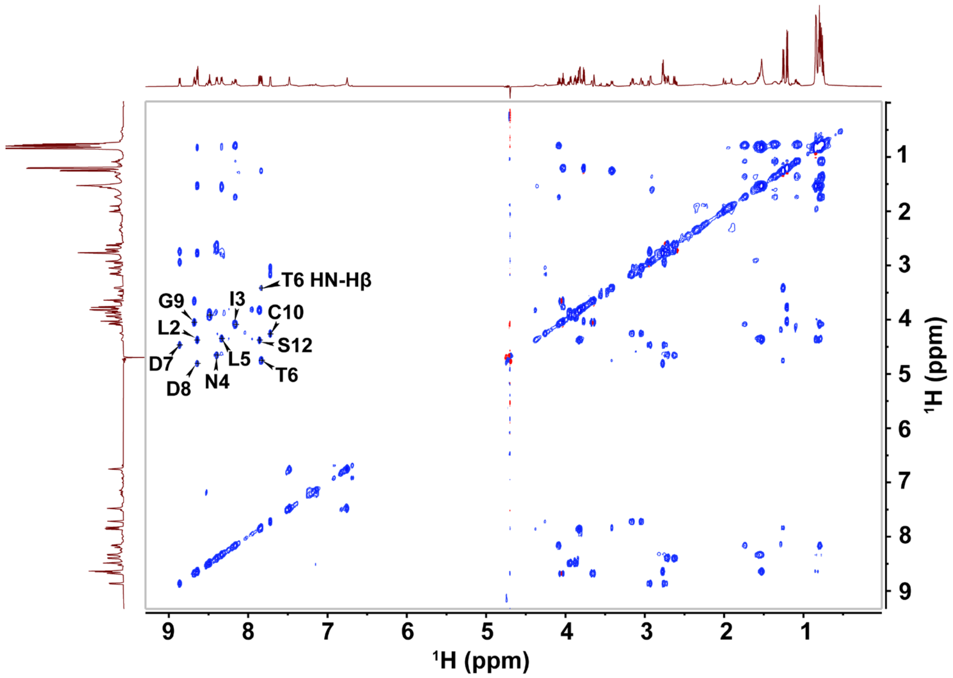

**Figure S11.** $^1$H-$^1$H TOCSY spectrum of LysC-digested mCoiA1$_{1\text{-}48}$-A36K(BCS$_A$) in 90% $H_2O$ and 10% $D_2O$. The amide proton assignments for each residue are annotated in the figure.

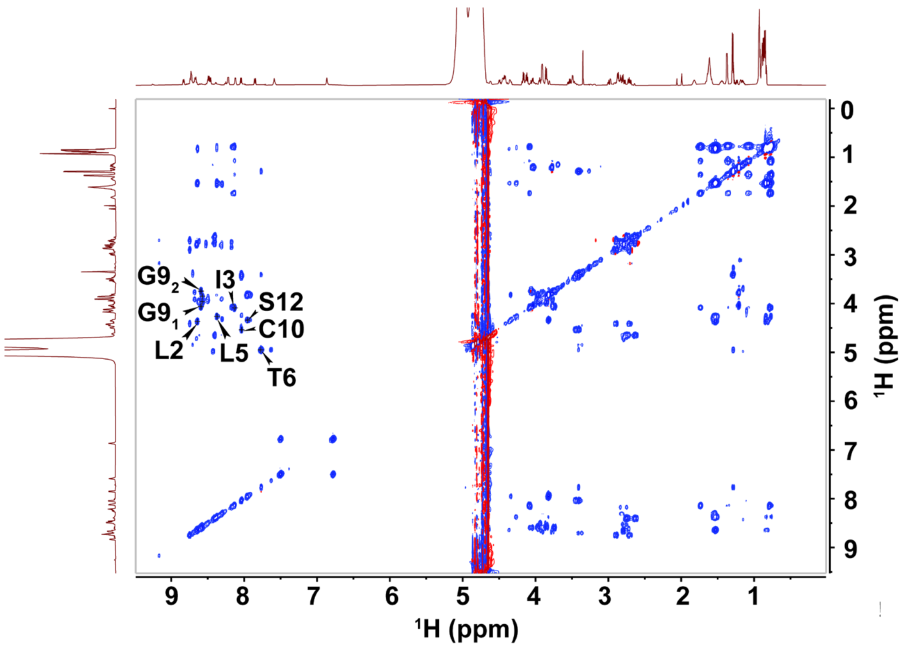

**Figure S12.** $^1$H-$^1$H TOCSY spectrum of $H_2O_2$-oxidized mCoiA1$_{1\text{-}48}$-A36K(BCS$_A$)$_{LysC,}$ in 90% $H_2O$ and 10% $D_2O$. The amide proton assignments for each residue are annotated in the figure.

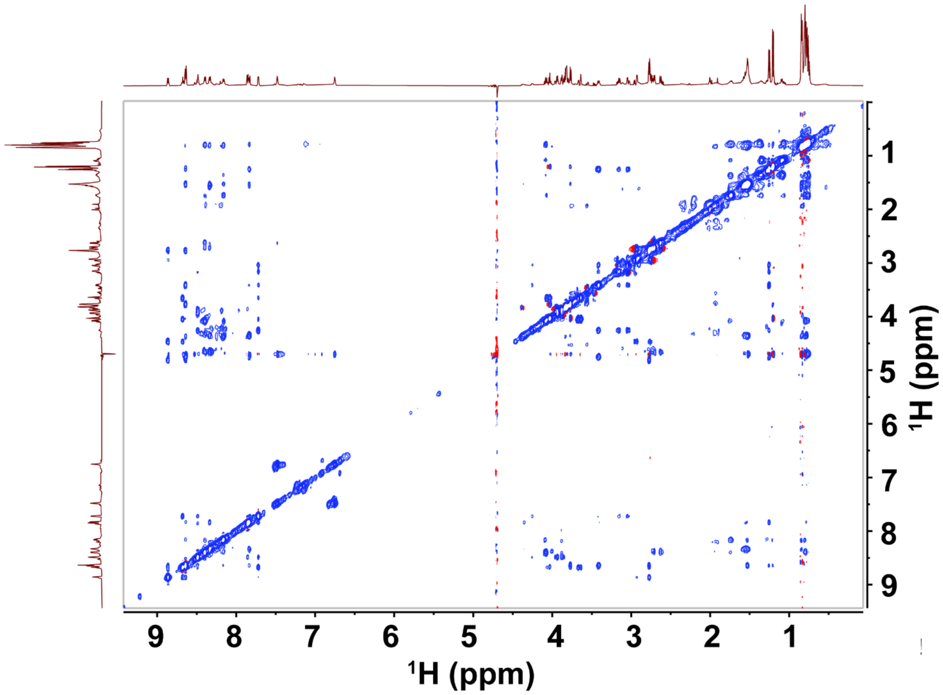

**Figure S13.** $^1$H-$^1$H NOESY spectrum of LysC-digested mCoiA1$_{1-48}$-A36K(BCS$_A$) in 90% $H_2O$ and 10% $D_2O$.

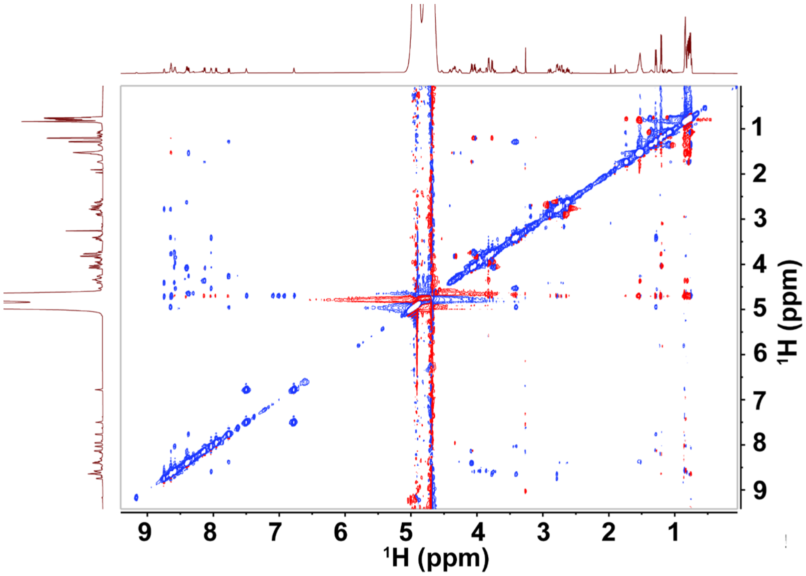

**Figure S14.** $^1$H-$^1$H NOESY spectrum of $H_2O_2$-oxidaized mCoiA1$_{1-48}$-A36K(BCS$_A$)$_{LysC,}$ in 90% $H_2O$ and 10% $D_2O$.

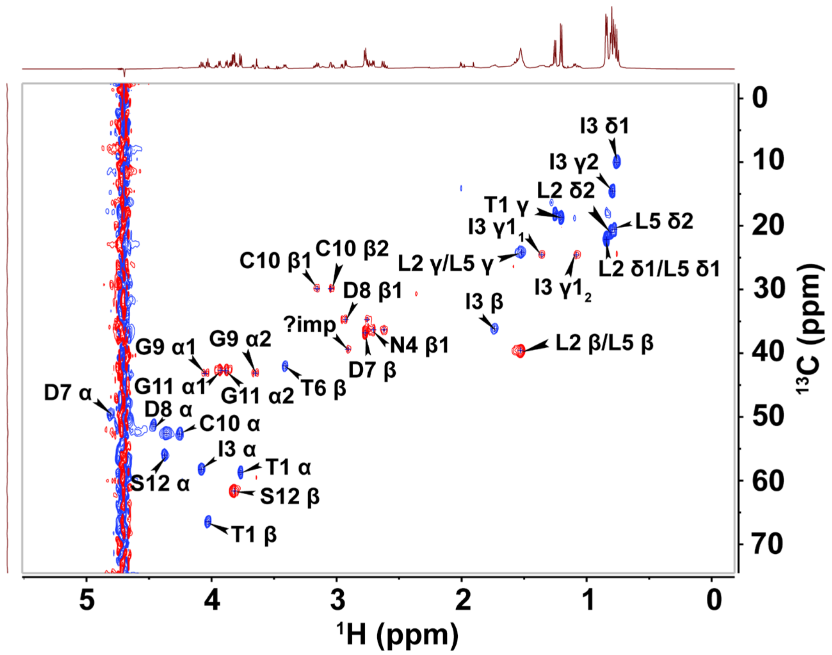

**Figure S15.** $^{1}H$-$^{13}C$ HSQC spectrum of LysC-digested mCoiA1$_{1\text{-}48}$-A36K(BCS$_A$) in 90% $H_2O$ and 10% $D_2O$. The cross-peaks are annotated in the figure, and their assignments are summarized in Table S6.

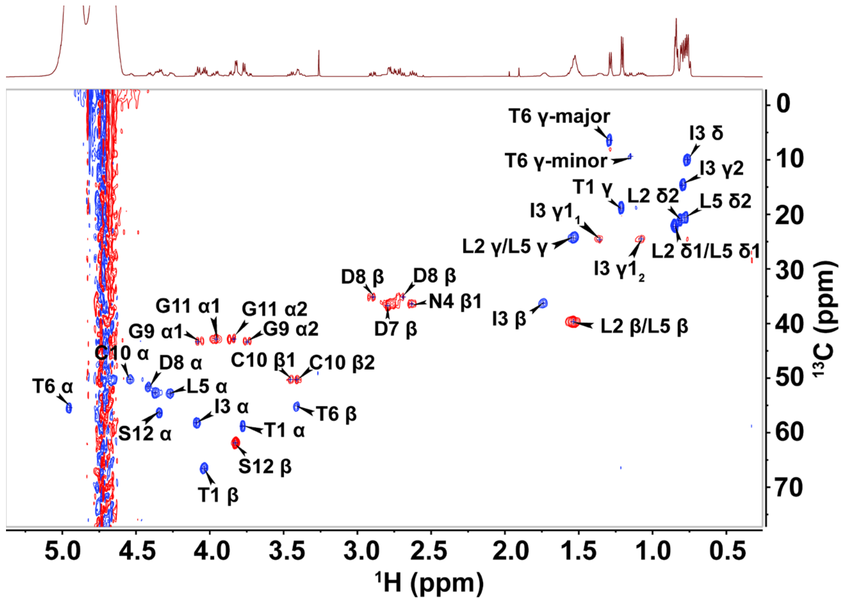

**Figure S16.** $^{1}H$-$^{13}C$ HSQC spectrum of $H_2O_2$-oxidized mCoiA1$_{1\text{-}48}$-A36K(BCS$_A$)$_{LysC}$ in 90% $H_2O$ and 10% $D_2O$. The cross-peak assignments are annotated in the figure. The cross-peaks are annotated in the figure, and their assignments are summarized in Table S6.

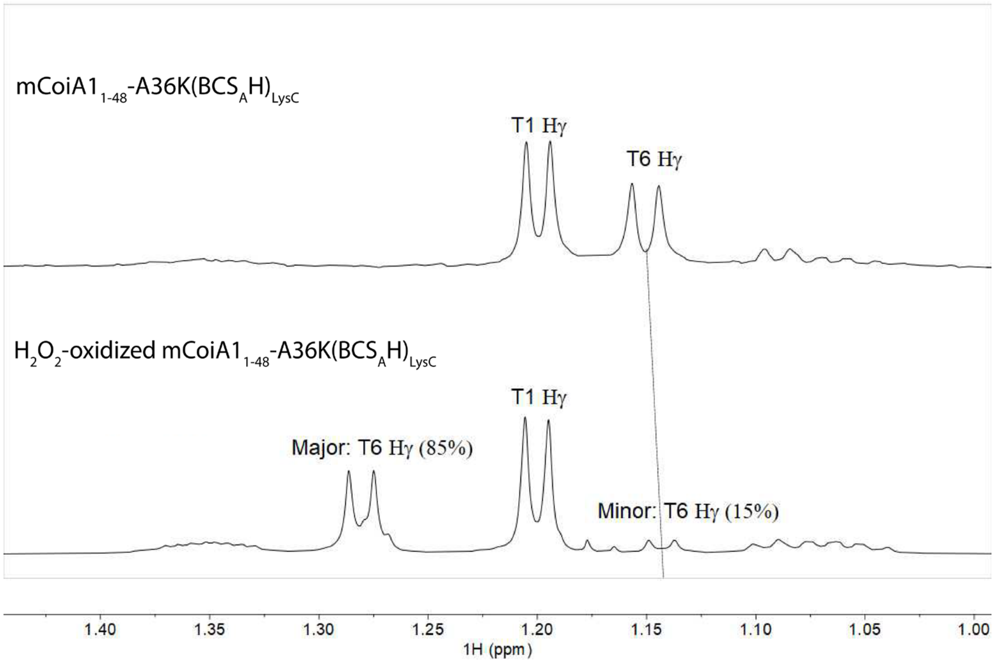

**Figure S17.** 1D $^1$H NMR spectra of mCoiA1$_{1\text{-}48}$-A36K(BCS$_A$)$_{LysC}$ and $H_2O_2$-oxidized mCoiA1$_{1\text{-}48}$-A36K(BCS$_A$)$_{LysC}$ showing the Hγ signals of former Thr6. Chemical oxidation with $H_2O_2$ yielded two diastereomers in an approximately 85:15 ratio (major:minor). The Thr6 Hγ signal of the CoiH-modified peptide closely matches that of the minor diastereomer of chemically oxidized peptide. This observation shows that CoiH is highly stereoselective.

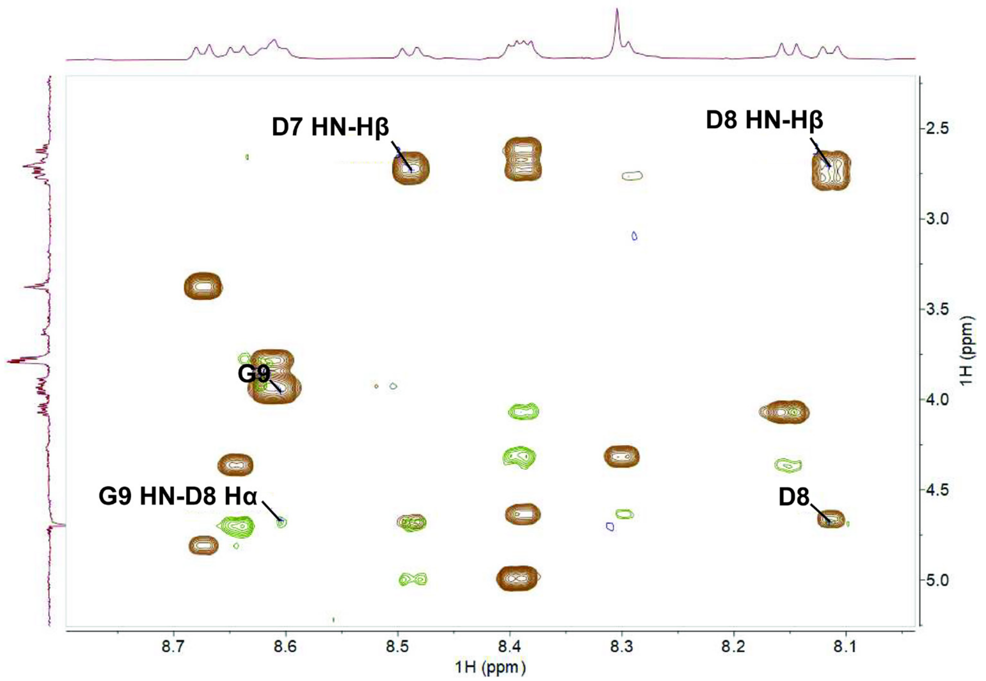

**Figure S18.** Overlay of $^1$H-$^1$H TOCSY and NOESY amide regions of LysC-digested mCoiA1$_{1-48}$-A36K(BCS$_A$H) in 90% $H_2O$ and 10% $D_2O$. The observed cross-peak between HN of Gly9 and Hα of Asp8, rather than between HN of Gly9 and Hβ of Asp8, confirms that Asp8 is a standard L-aspartate residue rather than an isoaspartate residue.

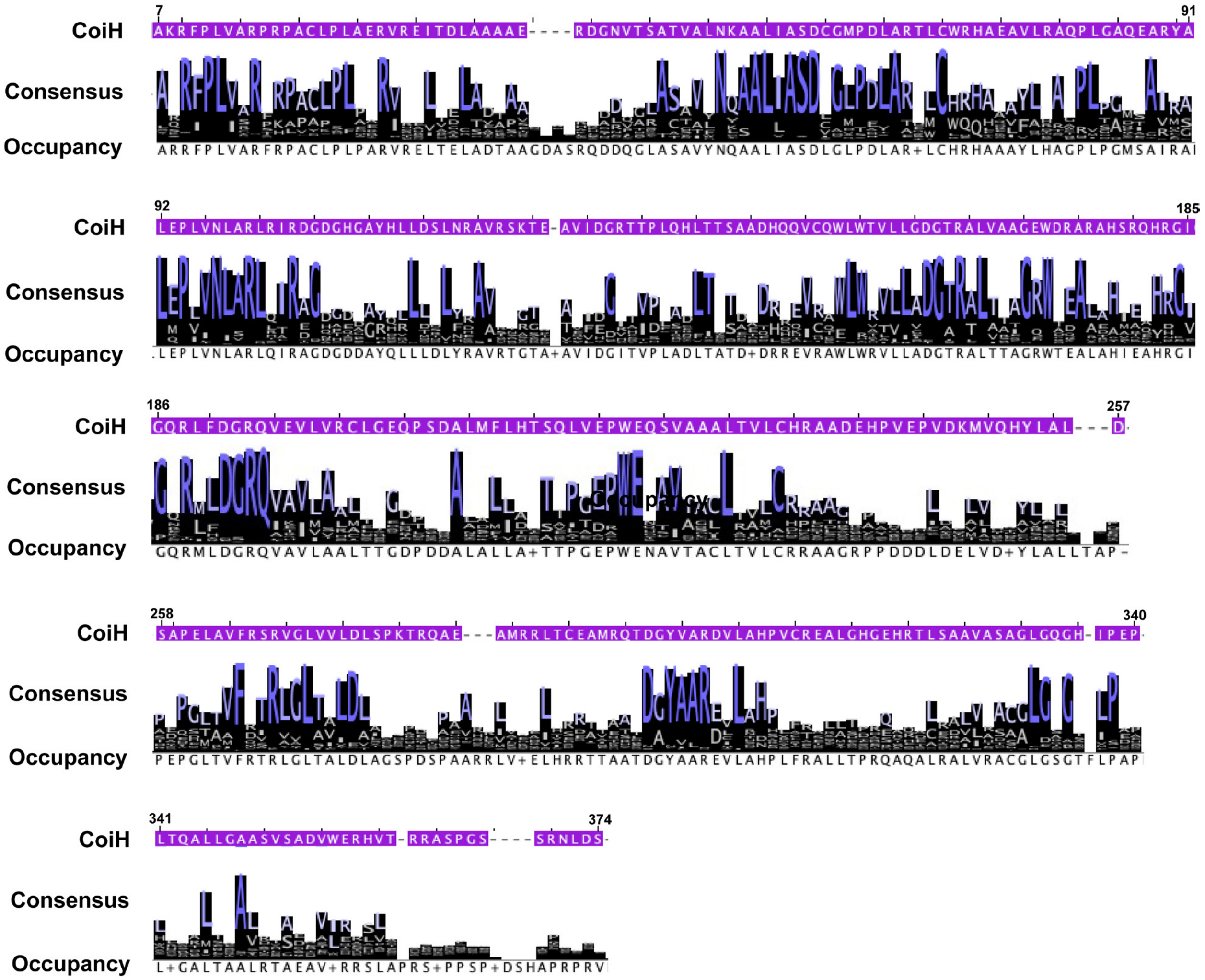

**Figure S19.** MAFFT alignment of CoiH and its homologs, using 4 to10 proteins selected from each cluster of the SSN analysis of CoiH and its homologs (Figure 6A); in total 133 proteins including CoiH, KitH, LydH, and PauH were aligned.

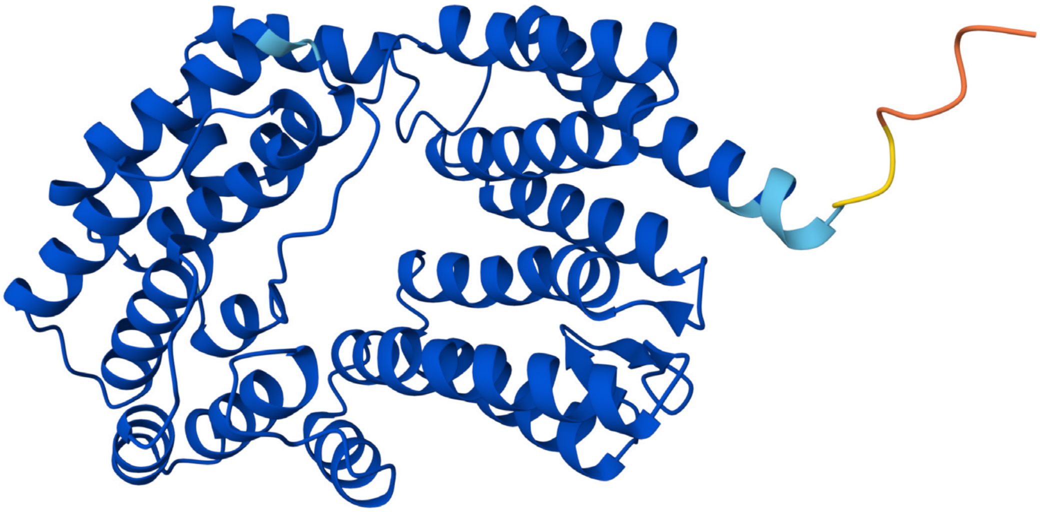

**Figure S20.** Overall AlphaFold 3-predicted structure of CoiH.

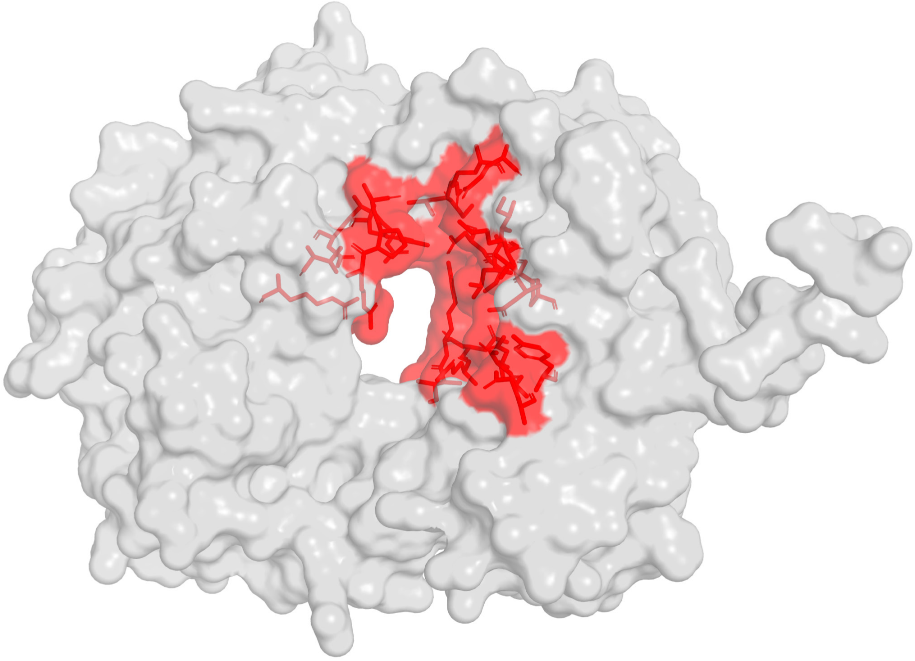

**Figure S21.** Surface representation of the AlphaFold 3-predicted structure of CoiH. The highest-ranked binding pocket identified by PrankWeb is highlighted in red, with the corresponding pocket residues shown as red sticks. Conserved residues surrounding the pocket (R26, N51, S58, D59, N97, R100, R268, and L332) were selected for site-directed mutagenesis to evaluate potential roles in catalysis.

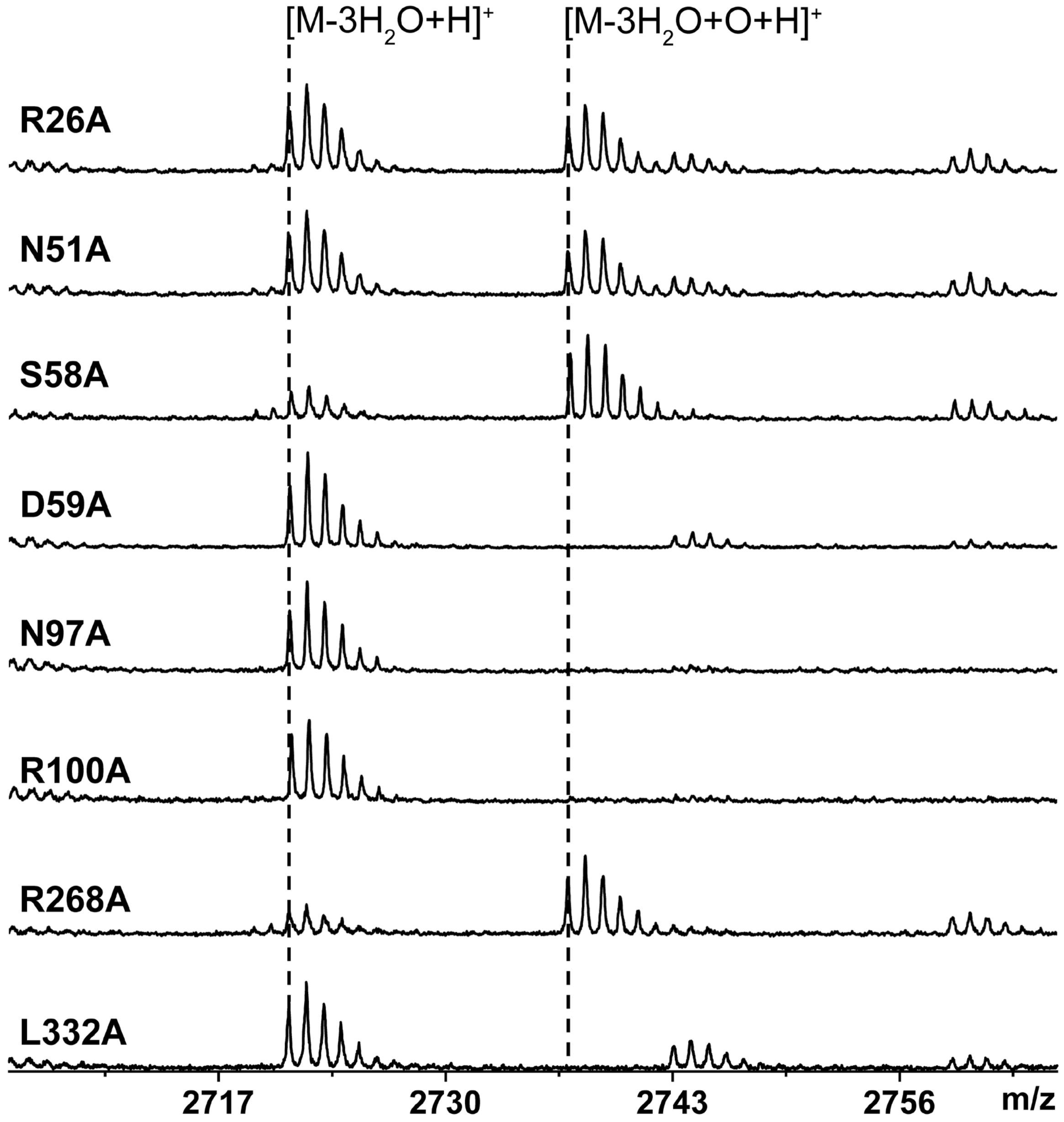

**Figure S22.** Conserved residues surrounding the highest-ranked pocket (Figure S21) were selected for site-directed mutagenesis to evaluate their potential roles in catalysis. CoiH variants (CoiH*) were coexpressed with SUMO-CoiA1BCS$_A$. mCoiA1(BCS$_A$H*) was digested by GluC and subsequently analyzed by MALDI-TOF MS. For calculated and observed m/z values, see Table S11.

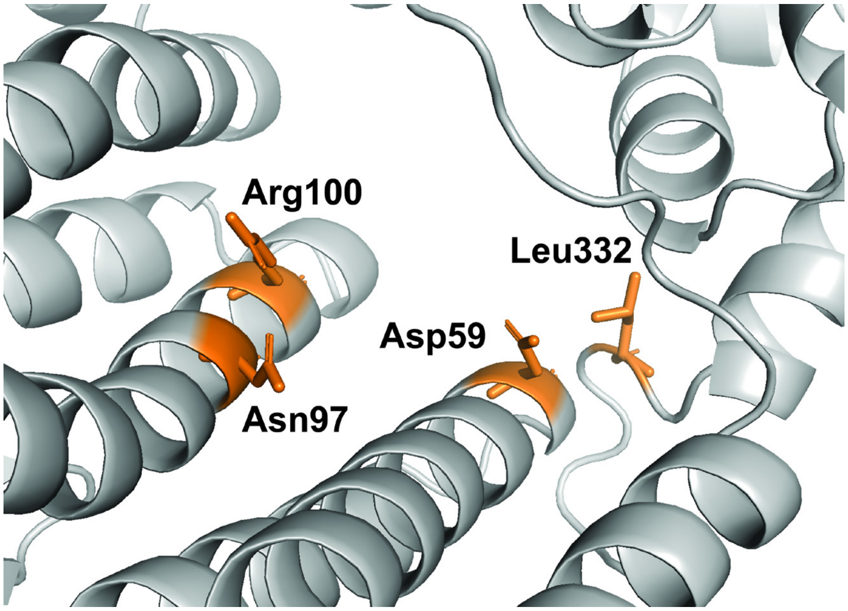

**Figure S23.** Four residues essential for sulfoxidation activity and/or enzyme stability in CoiH were identified by site-directed mutagenesis (Figure S22).

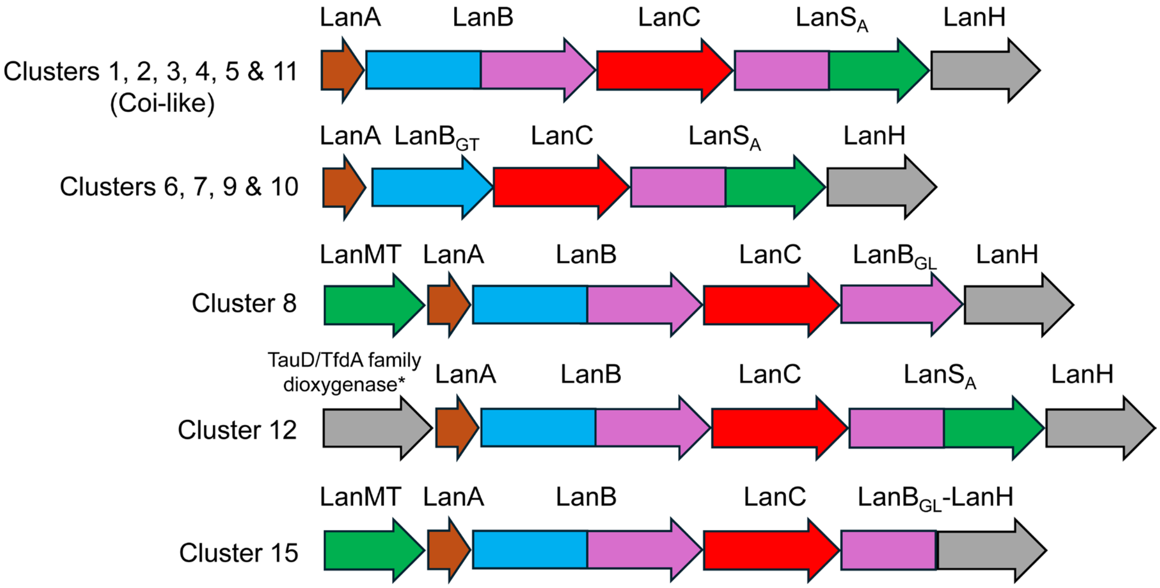

**Figure S24.** Representative BGC architectures of clusters that encode CoiH homologs identified in the SSN (Figure 6A, main text). LanMT, methyltransferase. LanB$_{GT}$, glutamyl transferase (blue). LanB$_{GL}$, glutamyl lyase (magenta). *Most BGCs of cluster 12 encode a TauD/TfdA family dioxygenase, but the KitH BGC does not have this gene. BGCs associated with clusters 6,7,9 and 10 lack a gene encoding LanBs with a canonical GL domain. Thus, Thr residues in the substrates of these BGCs are converted exclusively to (*E*)-Dhb.

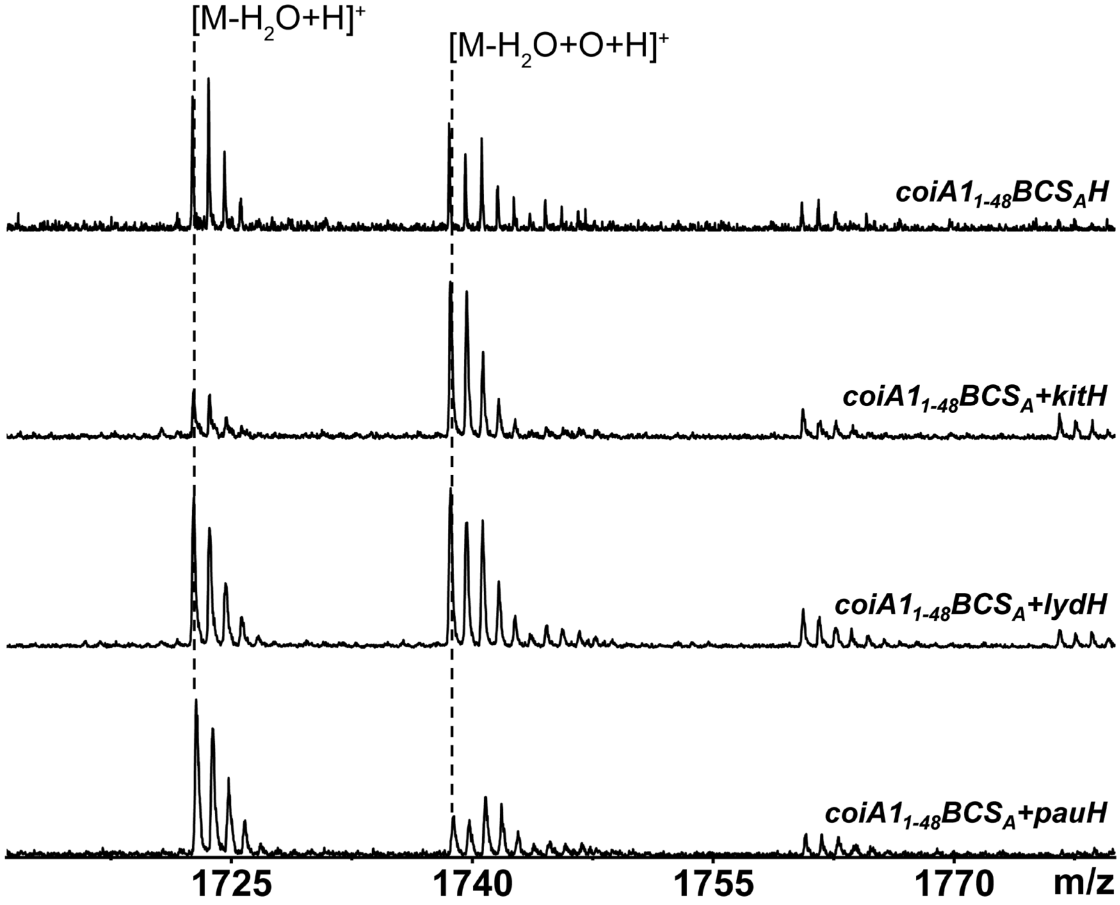

**Figure S25.** MALDI-TOF MS analysis of mCoiA1$_{1\text{-}48}$(BCS$_A$) modified by LanH (CoiH, KitH, LydH, and PauH) in *E. coli*. His$_6$-SUMO-CoiA1$_{1\text{-}48}$ was co-expressed with CoiB, CoiC, CoiS$_A$ and LanH, followed by GluC digestion. For calculated and observed m/z values, see Table S12.

**Table S1.** MALDI-TOF and ESI-MS/MS analysis of CoiA1 co-expressed with $CoiBCS_A$ or $CoiBCS_AH$ in *E. coli* and digested with GluC, corresponding to Figure 2.

| Ion Type | Charge State | Calculated m/z | Observed m/z | Mass Error (Δmass) |
|---|---|---|---|---|
| MALDI-TOF: $mCoiA1(BCS_A)_{GluC}$ | | | | |
| $M\text{-}3H_2O\text{+}H$ | +1 | 2721.076 | 2721.176 | 0.10 |
| $M\text{-}3H_2O\text{+}Na$ | +1 | 2743.066 | 2743.149 | 0.08 |
| $M\text{-}3H_2O\text{+}K$ | +1 | 2759.174 | 2759.125 | -0.05 |
| MALDI-TOF: $mCoiA1(BCS_AH)_{GluC}$ | | | | |
| $M\text{-}3H_2O\text{+}H$ | +1 | 2721.076 | 2721.208 | 0.13 |
| $M\text{-}3H_2O\text{+}16\text{+}H$ | +1 | 2737.076 | 2737.207 | 0.13 |
| $M\text{-}3H_2O\text{+}Na$ | +1 | 2743.066 | 2743.170 | 0.10 |
| $M\text{-}3H_2O\text{+}K$ | +1 | 2759.174 | 2759.167 | -0.01 |
| $M\text{-}3H_2O\text{+}16\text{+}Na$ | +1 | 2759.066 | 2759.167 | 0.10 |
| $M\text{-}3H_2O\text{+}16\text{+}K$ | +1 | 2775.174 | 2775.152 | -0.02 |
| Ion Type | Charge State | Calculated m/z | Observed m/z | Mass Error (Δppm) |
| ESI-MS/MS: $mCoiA1(BCS_A)_{GluC}$ | | | | |
| $b_4$ | +1 | 375.11470 | 375.1119 | -7.46 |
| $b_5$ | +1 | 462.14673 | 462.14414 | -5.60 |
| $b_6$ | +1 | 533.18384 | 533.1818 | -3.83 |
| $b_7$ | +1 | 634.23152 | 634.22876 | -4.35 |
| $b_8$ | +1 | 747.31558 | 747.31279 | -3.73 |
| $b_9$ | +1 | 860.39965 | 860.39714 | -2.92 |
| $b_{10}$ | +1 | 974.44257 | 974.44128 | -1.32 |
| $b_{11}$ | +1 | 1087.52664 | 1087.52315 | -3.21 |
| $b_{16}$ | +1 | 1560.64385 | 1560.64243 | -0.91 |
| $b_{18}$ | +1 | 1704.69734 | 1704.69847 | 0.66 |
| $b_{26}$ | +1 | 2418.93500 | 2418.93011 | -2.02 |
| $b_{26}$ | +2 | 1209.97116 | 1209.9741 | 2.43 |
| $b_{27}$ | +2 | 1266.99263 | 1266.99484 | 1.74 |
| $b_{28}$ | +2 | 1316.52683 | 1316.53037 | 2.69 |
| $y_2$ | +1 | 189.12341 | 189.12239 | -5.39 |
| $y_3$ | +1 | 303.16634 | 303.16507 | -4.19 |
| $y_{11}$ | +1 | 1017.40400 | 1017.40857 | 4.49 |
| $y_{12}$ | +1 | 1104.43603 | 1104.44023 | 3.80 |
| $y_{13}$ | +1 | 1161.45749 | 1161.4616 | 3.54 |
| $y_{18}$ | +1 | 1634.57471 | 1634.57817 | 2.12 |
| $y_{19}$ | +1 | 1747.65877 | 1747.66117 | 1.37 |
| $y_{20}$ | +1 | 1861.70170 | 1861.70701 | 2.85 |
| $y_{21}$ | +1 | 1974.78576 | 1974.78839 | 1.33 |

| | | | | |
|---|---|---|---|---|
| $y_{22}$ | +1 | 2087.86982 | 2087.86848 | -0.64 |
| $y_{23}$ | +1 | 2188.91750 | 2188.91882 | 0.60 |
| $y_{28}$ | +2 | 1317.53466 | 1317.53318 | -1.12 |
| M+2H | +2 | 1361.05067 | 1361.05419 | 2.59 |
| ESI-MS/MS: mCoiA1($BCS_AH$)$_{GluC}$ | | | | |
| $b_4$ | +1 | 375.11470 | 375.1141 | -1.60 |
| $b_5$ | +1 | 462.14673 | 462.14488 | -4.00 |
| $b_6$ | +1 | 533.18384 | 533.18067 | -5.95 |
| $b_7$ | +1 | 634.23152 | 634.22845 | -4.84 |
| $b_8$ | +1 | 747.31558 | 747.3122 | -4.52 |
| $b_9$ | +1 | 860.39965 | 860.39598 | -4.27 |
| $b_{10}$ | +1 | 974.44257 | 974.43807 | -4.62 |
| $b_{11}$ | +1 | 1087.52664 | 1087.5213 | -4.91 |
| $b_{18}$ | +1 | 1720.69634 | 1720.68333 | -7.56 |
| $b_{26}$ | +1 | 2434.93400 | 2434.93475 | 0.31 |
| $b_{26}$ | +2 | 1217.97066 | 1217.97171 | 0.86 |
| $b_{27}$ | +2 | 1274.99213 | 1274.99129 | -0.66 |
| $b_{28}$ | +2 | 1324.52633 | 1324.5259 | -0.32 |
| $y_2$ | +1 | 189.12341 | 189.12197 | -7.61 |
| $y_3$ | +1 | 303.16634 | 303.16515 | -3.93 |
| $y_{11}$ | +1 | 1017.40400 | 1017.40645 | 2.41 |
| $y_{18}$ | +1 | 1650.57371 | 1650.57484 | 0.68 |
| $y_{19}$ | +1 | 1763.65777 | 1763.66 | 1.26 |
| $y_{20}$ | +1 | 1877.70070 | 1877.70015 | -0.29 |
| $y_{21}$ | +1 | 1990.78476 | 1990.78526 | 0.25 |
| $y_{22}$ | +1 | 2103.86882 | 2103.85834 | -4.98 |
| $y_{28}$ | +2 | 1325.53416 | 1325.53064 | -2.66 |
| M+2H | +2 | 1369.05017 | 1369.04971 | -0.34 |

**Table S2.** MALDI-TOF analysis of CoiA1 variants co-expressed with CoiBCS$_A$H in *E. coli* and digested by TEV protease, corresponding to Figure 3.

| Ion Type | Charge State | Calculated m/z | Observed m/z | Mass Error (Δmass) |
|---|---|---|---|---|
| mCoiA1-D43A($BCS_AH$)$_{TEV}$ | | | | |
| M-3$H_2O$+H | +1 | 5729.563 | 5729.641 | 0.08 |
| M-3$H_2O$+16+H | +1 | 5745.563 | 5745.691 | 0.13 |
| mCoiA1-D44A($BCS_AH$)$_{TEV}$ | | | | |
| M-3$H_2O$+H | +1 | 5729.563 | 5729.695 | 0.13 |
| M-3$H_2O$+16+H | +1 | 5745.563 | 5745.767 | 0.20 |
| mCoiA1-D43A/D44A($BCS_AH$)$_{TEV}$ | | | | |
| M-3$H_2O$+H | +1 | 5685.573 | 5685.618 | 0.05 |

| | | | | |
|---|---|---|---|---|
| M-3$H_2O$+16+H | +1 | 5701.573 | 5701.609 | 0.04 |
| M-2$H_2O$+16+H | +1 | 5719.583 | 5719.644 | 0.06 |
| mCoiA1$_{1\text{-}48}$($BS_A$)$_{TEV}$ | | | | |
| M-$H_2O$+H | +1 | 4775.158 | 4775.240 | 0.08 |
| mCoiA1$_{1\text{-}48}$($BS_A$H)$_{TEV}$ | | | | |
| M-$H_2O$+H | +1 | 4775.158 | 4774.759 | -0.40 |
| mCoiA1$_{1\text{-}48}$($BCS_A$H)$_{TEV}$ | | | | |
| M-$H_2O$+H | +1 | 4775.158 | 4774.972 | -0.19 |
| M-$H_2O$+16+H | +1 | 4791.158 | 4790.985 | -0.17 |

**Table S3.** MALDI-TOF MS and ESI-MS/MS analysis of $H_2O_2$-treated mCoiA1$_{1\text{-}48}$($BCS_A$H)$_{TEV}$, mCoiA1$_{1\text{-}48}$-A36K($BCS_A$H)$_{LysC}$ and mCoiA1$_{1\text{-}48}$($BCS_A$H)$_{AspN}$, corresponding to Figure 4.

| Ion Type | Charge State | Calculated m/z | Observed m/z | Mass Error (Δmass) |
|---|---|---|---|---|
| MALDI-TOF: mCoiA1$_{1\text{-}48}$($BCS_A$H)$_{TEV}$ | | | | |
| M-$H_2O$+H | +1 | 4775.158 | 4774.972 | -0.19 |
| M-$H_2O$+16+H | +1 | 4791.158 | 4790.985 | -0.17 |
| MALDI-TOF: mCoiA1$_{1\text{-}48}$($BCS_A$H)$_{TEV}$+$H_2O_2$ | | | | |
| M-$H_2O$+2O+H | +1 | 4807.158 | 4806.349 | -0.81 |
| M-$H_2O$+16+O+H | +1 | 4807.158 | 4806.349 | -0.81 |
| MALDI-TOF: mCoiA1$_{1\text{-}48}$-A36K($BCS_A$)$_{LysC}$ | | | | |
| M-$H_2O$+H | +1 | 1190.536 | 1190.502 | -0.03 |
| MALDI-TOF: $H_2O_2$-oxidized mCoiA1$_{1\text{-}48}$-A36K($BCS_A$)$_{LysC}$ | | | | |
| M-$H_2O$+16+H | +1 | 1206.531 | 1206.469 | -0.06 |
| MALDI-TOF: mCoiA1$_{1\text{-}48}$-A36K($BCS_A$H)$_{LysC}$ | | | | |
| M-$H_2O$+16+H | +1 | 1206.531 | 1206.840 | 0.31 |
| Ion Type | Charge State | Calculated m/z | Observed m/z | Mass Error (Δppm) |
| ESI-MS/MS: $H_2O_2$-oxidized mCoiA1$_{1\text{-}48}$($BCS_A$H)$_{TEV}$ | | | | |
| $b_{26}$ | +3 | 843.38860 | 843.3848 | -4.51 |
| $b_{30}$ | +3 | 980.81146 | 980.8099 | -1.59 |
| $b_{31}$ | +3 | 1023.82566 | 1023.8246 | -1.04 |
| $b_{38}$ | +3 | 1234.90039 | 1234.9012 | 0.66 |
| $b_{39}$ | +3 | 1272.59508 | 1272.5936 | -1.16 |
| $b_{40}$ | +3 | 1310.28977 | 1310.2876 | -1.66 |
| $y_9$ | +1 | 879.31458 | 879.3141 | -0.55 |
| $y_{10}$ | +1 | 992.39865 | 992.3941 | -4.58 |
| $y_{11}$ | +1 | 1105.48271 | 1105.4807 | -1.82 |
| $y_{12}$ | +1 | 1206.53039 | 1206.5256 | -3.97 |
| ESI-MS/MS: mCoiA1$_{1\text{-}48}$($BCS_A$H)$_{AspN}$ | | | | |
| $b_2$ | +1 | 173.05573 | 173.0555 | -1.33 |

| | | | | |
|---|---|---|---|---|
| $b_3$ | +1 | 260.08776 | 260.087 | -2.92 |
| $b_3$-$H_2O$ | +1 | 242.07276 | 242.0764 | 15.04 |
| $b_4$ | +1 | 331.12487 | 331.1242 | -2.02 |
| $b_4$-$H_2O$ | +1 | 313.10987 | 313.1141 | 13.51 |
| $b_5$ | +1 | 432.17255 | 432.1713 | -2.89 |
| $b_5$-$H_2O$ | +1 | 414.15755 | 414.1614 | 9.30 |
| $b_6$ | +1 | 545.25661 | 545.2565 | -0.20 |
| $b_6$-$H_2O$ | +1 | 527.24161 | 527.2452 | 6.81 |
| $b_7$ | +1 | 658.34068 | 658.3413 | 0.94 |
| $b_7$-$H_2O$ | +1 | 640.32568 | 640.3284 | 4.25 |
| $b_8$-$H_2O$ | +1 | 754.3686 | 754.373 | 5.83 |
| $b_9$ | +1 | 885.46767 | 885.4662 | -1.66 |
| $b_9$-$H_2O$ | +1 | 867.45267 | 867.4542 | 1.76 |
| $y_1$ | +1 | 106.04991 | 106.0492 | -6.69 |
| $y_2$ | +1 | 163.07138 | 163.0716 | 1.35 |
| $y_2$-$H_2O$ | +1 | 145.05638 | 145.061 | 31.85 |
| $y_3$-$H_2O$ | +1 | 264.06456 | 264.0637 | -3.26 |
| $y_4$-$H_2O$ | +1 | 321.08603 | 321.0832 | -8.81 |
| $y_5$-$H_2O$ | +1 | 436.11297 | 436.1122 | -1.77 |
| $y_6$-$H_2O$ | +1 | 551.13991 | 551.141 | 1.98 |
| $y_7$-$H_2O$ | +1 | 634.17259 | 634.1767 | 6.48 |
| $y_8$ | +1 | 765.27165 | 765.2703 | -1.76 |
| $y_8$-$H_2O$ | +1 | 747.25665 | 747.2585 | 2.48 |
| $y_9$ | +1 | 879.31458 | 879.3133 | -1.46 |
| $y_9$-$H_2O$ | +1 | 861.29958 | 861.3033 | 4.32 |
| $y_{10}$ | +1 | 992.39865 | 992.3974 | -1.26 |
| $y_{10}$-$H_2O$ | +1 | 974.38365 | 974.3849 | 1.28 |
| $y_{11}$ | +1 | 1105.48271 | 1105.4814 | -1.19 |
| $y_{11}$-$H_2O$ | +1 | 1087.46771 | 1087.466 | -1.57 |

**Table S4.** ESI-MS/MS analysis of mCoiA$1_{1\text{-}48}$-A36K$(BCS_A)_{LysC}$ and $H_2O_2$-oxidized mCoiA$1_{1\text{-}48}$-A36K$(BCS_A)_{LysC}$, corresponding to Figures S4-S5.

| Ion Type | Charge State | Calculated m/z | Observed m/z |
|---|---|---|---|
| mCoiA$1_{1\text{-}48}$-A36K$(BCS_A)_{LysC}$ | | | |
| $b_2$ | +1 | 197.13 | 197.13 |
| $b_3$ | +1 | 310.21 | 310.21 |
| $b_4$ | +1 | 424.26 | 424.25 |
| $b_5$ | +1 | 537.34 | 537.33 |
| $b_{10}$ | +1 | 1010.46 | 1010.45 |
| $b_{11}$ | +1 | 1089.49 | 1089.52 |
| $y_7$ | +1 | 636.19 | 636.19 |

| | | | |
|---|---|---|---|
| $y_8$ | +1 | 749.27 | 749.28 |
| $y_9$ | +1 | 863.32 | 863.32 |
| $y_{10}$ | +1 | 976.40 | 976.40 |
| $H_2O_2$-oxidized mCoiA1$_{1\text{-}48}$-A36K(BCS$_{A}$)$_{LysC}$ | | | |
| $b_2$ | +1 | 197.13 | 197.12 |
| $b_3$ | +1 | 310.21 | 310.21 |
| $b_4$ | +1 | 424.26 | 424.25 |
| $b_5$ | +1 | 537.34 | 537.33 |
| $y_8$ | +1 | 781.28 | 781.27 |
| $y_9$ | +1 | 895.32 | 895.31 |
| $y_{10}$ | +1 | 1008.40 | 1008.41 |

**Table S5.** $^{1}H$ and $^{13}C$ chemical shifts of LysC-digested mCoiA1$_{1\text{-}48}$-A36K(BCS$_A$H) in 90% $H_2O$ and 10% $D_2O$ at 25 °C. *

| # | AA ID | N-H | αH/C | βH/C | γH/C | other |
|---|---|---|---|---|---|---|
| 1 | T | - | 3.77<br>58.7 | 4.03<br>66.5 | 1.21<br>18.7 | |
| 2 | L | 8.64 | 4.37<br>52.6 | 1.56<br>39.6 | 1.53<br>24.2 | 0.84, 22.2<br>0.81, 21.0 |
| 3 | I | 8.15 | 4.08<br>58.2 | 1.74 (CH)<br>36.2 | γ1($CH_2$):<br>1.36,1.10<br>24.6 | γ2($CH_3$): 0.79<br>14.6<br>δ1($CH_3$): 0.76<br>(t)<br>10.1 |
| 4 | N | 8.39 | 4.64<br>52.6 | 2.73, 2.62<br>36.4 | | 7.50, 6.77 |
| 5 | L | 8.30 | 4.32<br>52.6 | 1.55<br>39.6 | 1.53<br>24.2 | 0.84, 22.2<br>0.78, 20.6 |
| **6** | **(T)<br>Abu** | **8.40** | **4.99<br>53.0** | **3.62<br>60.2 (-S=O)**<br>(in 100%$D_2O$: 3.74, 60.6) | **1.16<br>9.2<br>(-S=O)** | |
| 7 | D | 8.49 | 4.68<br>50.4** | 2.74, 2.70<br>36.0 | | |
| **8** | **D** | **8.11** | **4.66<br>50.4**** | **2.78, 2.69<br>36.0** | | **L-aspartate** |
| 9 | G | 8.596 | 3.94<br>42.8 | | | |
| **10** | **(C)<br>Ala** | **8.674** | **4.81<br>50.6**** | **3.38<br>48.2 (-S=O)**<br>(in 100% $D_2O$: 3.44, 3.37, 48.3) | | |
| 11 | G | 8.609 | 3.90, 3.79<br>43.9 | | | |
| 12 | S | 7.84 | 4.27<br>57.0 | 3.79<br>62.1 | | |

* Chemical shifts that are diagnostic for sulfoxides are in red font and are either downfield shifted (Cβ) or upfield shifted (Cγ) compared to the methyllanthionine in the CoiBCS$_A$-modified peptide (Table S6). Thr6 residue converted to MeLan is shown as (T) Abu. Cys10 residue converted to MeLan is shown as (C) Ala. $^{13}C$ chemical shifts were measured for the sample in 100% $D_2O$.

**Table S6** $^{1}H$ and $^{13}C$ chemical shifts of mCoiA1$_{1\text{-}48}$-A36K(BCS$_A$)$_{LysC}$ and $H_2O_2$-oxidized mCoiA1$_{1\text{-}48}$-A36K(BCS$_A$)$_{LysC}$ in 90% $H_2O$ and 10% $D_2O$ at 25 °C.

| # | AA ID | N-H | αH/C | βH/C | γH/C | other |
|---|---|---|---|---|---|---|
| 1 | T | - | 3.77<br>58.7 | 4.03<br>66.5 | 1.21<br>18.7 | |
| | | - | 3.77<br>58.7 | 4.03<br>66.4 | 1.20<br>18.8 | |
| 2 | L | 8.64 | 4.37<br>52.6 | 1.56<br>39.6 | 1.53<br>24.2 | 0.84, 22.2<br>0.81, 21.0 |
| | | 8.64 | 4.37<br>52.6 | 1.53<br>39.6 | 1.53<br>24.2 | 0.846, 21.8<br>0.81, 21.0 |
| 3 | I | 8.16 | 4.08<br>58.2 | 1.74 (CH)<br>36.2 | γ1CH$_2$:<br>1.36,1.10<br>24.6 | γ2CH$_3$: 0.79<br>14.6<br>δ1CH$_3$: 0.76 (t)<br>10.1 |
| | | 8.13 | 4.08<br>58.1 | 1.73 (CH)<br>36.2 | γ1CH$_2$: 1.35,<br>1.07<br>24.5 | γ2CH$_3$: 0.79<br>14.5<br>δ1CH$_3$: 0.76 (t)<br>10.0 |
| 4 | N | 8.40 | 4.66<br>52.6 | 2.72, 2.63<br>36.4 | | δNH2: 7.48, 6.75 |
| | | 8.40 | 4.65<br>- | 2.75, 2.62<br>36.3 | | δNH2: 7.50, 6.78 |
| 5 | L | 8.34 | 4.35<br>52.6 | 1.55<br>39.6 | 1.53<br>24.2 | 0.84, 22.2<br>0.78, 20.6 |
| | | 8.37 | 4.267<br>52.8 | 1.53<br>39.6 | 1.53<br>24.2 | 0.838, 22.2<br>0.77, 20.6 |
| **6** | **(T)<br>Abu** | 7.834 | 4.75<br>53.0 | 3.42 (CH)<br>42.1 | 1.25<br>18.2 | |
| | | 7.76 | 4.95<br>55.4 | 3.40 (CH)<br>**55.2**<br>**(-S=O)** | **Major: 1.28**<br>**6.44 (-S=O);**<br>**minor: 1.14, 9.5**<br>**(-S=O)** | |
| 7 | D | 8.642 | 4.81<br>50.6 | 2.78<br>36.8 | | |
| | | 8.65 | 4.70<br>- | 2.79<br>36.7 | | |
| 8 | D | 8.863 | 4.47<br>50.4 | 2.94, 2.75<br>34.8 | **L-aspartate** | |

| | | | | | |
|---|---|---|---|---|---|
| | | 8.74 | 4.41<br>51.5 | 2.90, 2.71<br>35.0 | |
| 9 | G | 8.678 | 4.05, 3.65<br>43.1 | | |
| | | 8.59 | 4.06, 3.74<br>43.2 | | |
| **10** | **(C) Ala** | 7.72 | 4.26<br>50.4 | 3.17, 3.04<br>29.8 | NOE observed between T6 Hβ-C10 Hβ, T6 Hγ-C10 Hβ, etc |
| | | 8.03 | 4.535<br>50.1 | 3.44, 3.40<br>**50.2**<br>**(-S=O)** | NOE observed between T6 Hγ-C10 Hβ, etc |
| 11 | G | 8.49 | 3.94, 3.88<br>42.6 | | |
| | | 8.57 | 3.95, 3.84<br>42.8 | | |
| 12 | S | 7.86 | 4.38<br>55.8 | 3.83<br>61.7 | |
| | | 7.96 | 4.34<br>56.3 | 3.82<br>61.8 | |

* In each residue entry, mCoiA1$_{1\text{-}48}$(BCS$_{A}$)$_{LysC}$ is shown in the top row and $H_2O_2$-oxidized mCoiA1$_{1\text{-}48}$(BCS$_{A}$)$_{LysC}$ is shown in the bottom row. Chemical shifts that are diagnostic for sulfoxides are in red font and are either downfield shifted (Cβ) or upfield shifted (Cγ) compared to other methyllanthionines. Thr6 residue converted to MeLan is shown as (T) Abu. Cys10 residue converted to MeLan is shown as (C) Ala.

**Table S7.** MALDI-TOF MS analysis of HPLC-purified mCoiA1$_{1-48}$-A36K(BCS$_A$H) digested with LysC, corresponding to Figure 5A.

| Ion Type | Charge State | Calculated m/z | Observed m/z | Mass Error (Δmass) |
|---|---|---|---|---|
| M-$H_2O$+H | +1 | 1190.525 | 1190.556 | 0.03 |
| M-$H_2O$+16+H | +1 | 1206.525 | 1206.556 | 0.03 |

**Table S8.** Similarity of CoiH with other oxygenases involved in sulfoxidation for natural product biosynthesis. None of the enzymes have folds like the predicted fold for CoiH.

| **Enzyme** | | **GarO**[9] | **SolS**[10] | **UstF1**[11] | **EgtB**[12] | **OvoA**[13] |
|---|---|---|---|---|---|---|
| **CoiH** | Identity | 5.0% | 12.8% | 2.2% | 11.7% | 4.8% |
| | Similarity | 5.9% | 18.0% | 2.9% | 17.4% | 7.9% |

**Table S9.** Typical conserved motif in oxygenases that are not found in the LanH sequences.

| **Oxygenases** | **Typical Motifs** |
|---|---|
| Flavin-dependent oxygenases | GXGXXG (at N-terminus) |
| Non-heme iron oxygenase | HXD |
| | HXE |
| Cytochrome P450 | FXXGXRXCXG |
| | EXXR |
| Rieske Oxygenase | $CXHX_{15-17}CX_2H$ |
| | $DX_2HX_4H$ |
| α-Ketoglutarate-dependent oxygenase | HXD/E...H |
| Copper oxygenase | H-Xn-H |

**Table S10.** Conserved residues in CoiH and in its predicted ligand-binding pocket. Only residues with consensus higher than 90% are shown.

| Amino acid position in CoiH | Amino acid in CoiH | Consensus | Amino acids in predicted pocket |
|---|---|---|---|
| 11 | P | 92.4% | |
| 12 | L | 92.4% | |
| 15 | R | 90.1% | |
| 26 | R | 91.6% | R26 |
| 46 | A | 90.1% | |
| 51 | N | 97% | N51 |
| 53 | A | 96.9% | |
| 54 | L | 97.7% | |
| 57 | A | 96.2% | |
| 58 | S | 84.7% | S58 |
| 59 | D | 97.7% | D59 |
| 66 | A | 97.7% | |
| 70 | C | 97.7% | |
| 88 | A | 90.8% | |
| 92 | L | 100% | |
| 94 | P | 100% | |
| 97 | N | 100% | N97 |
| 100 | R | 100% | R100 |
| 101 | L | 96.2% | |
| 104 | R | 100% | R104 |
| 106 | G | 92.4% | |
| 115 | L | 97.7% | |
| 121 | A | 95.4% | |
| 154 | L | 96.9% | |
| 161 | D | 94.7% | |
| 162 | G | 99.2% | |
| 164 | R | 100% | |
| 166 | L | 100% | |
| 170 | G | 97.7% | |
| 172 | W | 99.2% | |
| 175 | A | 98.5% | |
| 186 | G | 90.1% | |
| 188 | R | 92.4% | |
| 191 | D | 94.7% | |
| 192 | G | 93.1% | |
| 193 | R | 100% | |
| 194 | Q | 100% | |
| 209 | A | 100% | |
| 222 | W | 98.5% | |

| | | | |
|---|---|---|---|
| 223 | E | 99.2% | |
| 230 | L | 99.2% | |
| 263 | F | 98.5% | |
| 268 | R | 95.4% | R268 |
| 271 | L | 96.2% | |
| 298 | D | 93.9% | |
| 300 | Y | 93.9% | Y300 |
| 303 | R | 95.4% | R303 |
| 306 | L | 96.9% | |
| 332 | L | 94.7% | L332 |
| 348 | A | 91.6% | |

**Table S11.** MALDI-TOF MS analysis of GluC-digested CoiA1 modified by CoiBCS$_A$ and mutant CoiH, corresponding to Figure S22.

| Mutant CoiH | Ion Type | Charge State | Calculated m/z | Observed m/z | Mass Error (Δmass) |
|---|---|---|---|---|---|
| R26A | M+H-3$H_2O$ | +1 | 2721.076 | 2721.042 | -0.03 |
| | M+H-3$H_2O$+O | +1 | 2737.076 | 2737.017 | -0.06 |
| N51A | M+H-3$H_2O$ | +1 | 2721.076 | 2721.086 | 0.01 |
| | M+H-3$H_2O$+O | +1 | 2737.076 | 2737.080 | 0.00 |
| S58A | M+H-3$H_2O$ | +1 | 2721.076 | 2721.149 | 0.07 |
| | M+H-3$H_2O$+O | +1 | 2737.076 | 2737.138 | 0.06 |
| D59A | M+H-3$H_2O$ | +1 | 2721.076 | 2721.087 | 0.01 |
| N97A | M+H-3$H_2O$ | +1 | 2721.076 | 2721.062 | -0.01 |
| R100A | M+H-3$H_2O$ | +1 | 2721.076 | 2721.175 | 0.10 |
| R268A | M+H-3$H_2O$ | +1 | 2721.076 | 2721.097 | 0.02 |
| | M+H-3$H_2O$+O | +1 | 2737.076 | 2737.062 | -0.01 |
| L332A | M+H-3$H_2O$ | +1 | 2721.076 | 2721.012 | -0.06 |

**Table S12.** MALDI-TOF MS analysis of GluC-digested CoiA1$_{1-48}$ modified by CoiBCS$_A$ and LanH (CoiH, KitH, LydH, or PauH), corresponding to Figure S25.

| LanH | Ion Type | Charge State | Calculated m/z | Observed m/z | Mass Error (Δmass) |
|---|---|---|---|---|---|
| CoiH | M+H-$H_2O$ | +1 | 1722.702 | 1722.578 | -0.12 |
| | M+H-$H_2O$+O | +1 | 1738.702 | 1738.566 | -0.14 |
| KitH | M+H-$H_2O$ | +1 | 1722.702 | 1722.633 | -0.07 |
| | M+H-$H_2O$+O | +1 | 1738.702 | 1738.645 | -0.06 |
| LydH | M+H-$H_2O$ | +1 | 1722.702 | 1722.640 | -0.06 |
| | M+H-$H_2O$+O | +1 | 1738.702 | 1738.640 | -0.06 |
| PauH | M+H-$H_2O$ | +1 | 1722.702 | 1722.826 | 0.12 |
| | M+H-$H_2O$+O | +1 | 1738.702 | 1738.825 | 0.12 |

**Table S13.** Plasmids used in this study.

| Plasmids | Genes | Resistance | Source |
|---|---|---|---|
| pRSFDuet-His6-SUMO-CoiA1-CoiB | *His6-SUMO-coiA1*, *coiB* | $Kan^r$ | [14] |
| pRSFDuet-His6-SUMO-CoiA1-D43A-CoiB | D43A in *His6-SUMO-coiA1, CoiB* | $Kan^r$ | This work |
| pRSFDuet-His6-SUMO-CoiA1-D44A-CoiB | D44A in *His6-SUMO-coiA1, CoiB* | $Kan^r$ | This work |
| pRSFDuet-His6-SUMO-CoiA1-D43A/D44A-CoiB | D43A&D44A in *His6-SUMO-coiA1, CoiB* | $Kan^r$ | This work |
| pRSFDuet-His6-SUMO-CoiA1$_{1\text{-}48}$-CoiB | *His6-SUMO-coiA1*$_{1\text{-}48}$, *coiB* | $Kan^r$ | This work |
| pRSFDuet-His6-SUMO-CoiA1$_{1\text{-}48}$-A36K-CoiB | *His6-SUMO-coiA1*$_{1\text{-}48}^{A36K}$, *coiB* | $Kan^r$ | This work |
| pETDuet-CoiC-CoiS$_A$ | *coiC*, *coiS*$_A$ | $Amp^r$ | [14] |
| pETDuet-CoiS$_A$ | *coiS*$_A$ | $Amp^r$ | This work |
| pCDFDuet-CoiH | *coiH* | $Str^r$ | Previous work |
| pETDuet-CoiC-CoiS$_A$-KitH | *coiC, coiS*$_A$*, kitH* | $Str^r$ | This work |
| pETDuet-CoiC-CoiS$_A$-LydH | *coiC, coiS*$_A$*, lydH* | $Str^r$ | This work |
| pETDuet-CoiC-CoiS$_A$-PauH | *coiC, coiS*$_A$*, pauH* | $Str^r$ | This work |
| pCDFDuet-CoiH-R26A | R26A in *coiH* | $Str^r$ | This work |
| pCDFDuet-CoiH-N51A | N51A in *coiH* | $Str^r$ | This work |
| pCDFDuet-CoiH-S58A | S58A in *coiH* | $Str^r$ | This work |
| pCDFDuet-CoiH-D59A | D59A in *coiH* | $Str^r$ | This work |
| pCDFDuet-CoiH-N97A | N97A in *coiH* | $Str^r$ | This work |
| pCDFDuet-CoiH-R100A | R100A in *coiH* | $Str^r$ | This work |
| pCDFDuet-CoiH-R268A | R268A in *coiH* | $Str^r$ | This work |
| pCDFDuet-CoiH-L332A | L332A in *coiH* | $Str^r$ | This work |

**Table S14.** Primers used in this study.

| Primer name | Sequence (5'-3') |
|---|---|
| D43A-F | TTGACGGCTGATGGATGTGGCTCCACTTGCAGC |
| D43A-R | TCCATCAGCCGTCAAGTTAATCAGGGTCGCAGAG |
| D44A-F | ACGGATGCTGGATGTGGCTCCACTTGCAGCAG |
| D44A-R | ACATCCAGCATCCGTCAAGTTAATCAGGGTCG |
| D43A/D44A-F | TGACGGCTGCTGGATGTGGCTCCACTTGCAGCAGC |
| D43A/D44A-R | ATCCAGCAGCCGTCAAGTTAATCAGGGTCGCAGAG |
| CoiA1$_{1-48}$-F | TAATGCGGCCGCATAATGCTTAAGTC |
| CoiA1$_{1-48}$-R | TATGCGGCCGCATTAGGAGCCACATCCATCATCCGTCAAG |
| CoiA1$_{1-48}$-A36K-F | CGGCTCTAAAACCCTGATTAACTTGACGGATGATG |
| CoiA1$_{1-48}$-A36K-R | AGGGTTTTAGAGCCGTCGTCACTTTCTAAAACTG |
| CoiS$_A$-F | AGAAGGAGATATACCTGCGGCCGCATAATGCTTAAGTCGAAC |
| CoiS$_A$-R | GGTATATCTCCTTCTTAAAGTTAAACAAAATTATTTCTAG |
| KitH-F | TTAGTAAAGTATAAGGGAGGAATTAAATGCATCCGACAGGTCCGATTGCACAG |
| KitH-R | GTGGCAGCAGCCTAGGTTAATTAGGGGGTGGAATTCAGAGGATGTG |
| LydH-F | TTAGTAAAGTATAAGGGAGGAATTAAATGGACGCGTCTCAGATCGTGCGTC |
| LydH-R | GTGGCAGCAGCCTAGGTTAATTATGCGGAAGCTGATGGCGCCGCAG |
| PauH-F | TTAGTAAAGTATAAGGGAGGAATTAAATGGTCGATCTCGATCGTATTGCCCGTC |
| PauH-R | GTGGCAGCAGCCTAGGTTAATTATACCGTCGCGGCCTCGCTCATAG |
| Vector-CS$_A$-F | TTAACCTAGGCTGCTGCCACCGCTG |
| Vector-CS$_A$-F | CCTCCCTTATACTTTACTAATTACTGCCATTCGATCACGATTCTG |
| CoiH-R26A-F | CGGAAGCCGTGCGGGAGATTACCGATCTGGCAG |
| CoiH-R26A-R | CCCGCACGGCTTCCGCGAGTGGCAGACACGCAGGGCGTG |
| CoiH-N51A-F | GCCCTGGCCAAAGCGGCACTTATCGCCTCTGATTG |
| CoiH-N51A-R | CGCTTTGGCCAGGGCAACGGTTGCTGAGGTGACG |
| CoiH-S58A-F | ATCGCCGCCGATTGCGGGATGCCAGATCTCGCTC |
| CoiH-S58A-R | GCAATCGGCGGCGATAAGTGCCGCTTTGTTCAGG |
| CoiH-D59A-F | GCCTCTGCCTGCGGGATGCCAGATCTCGCTCGCAC |
| CoiH-D59A-R | CCCGCAGGCAGAGGCGATAAGTGCCGCTTTGTTC |
| CoiH-N97A-F | CTGGTGGCCCTGGCGCGCTTGCGTATCCGCGATG |
| CoiH-N97A-R | CGCCAGGGCCACCAGCGGTTCCAACGCGTAACG |
| CoiH-R100A-F | CTGGCGGCCTTGCGTATCCGCGATGGCGACGGAC |
| CoiH-R100A-R | ACGCAAGGCCGCCAGATTCACCAGCGGTTCCAAC |
| CoiH-R268A-F | CGTTCCGCCGTTGGCTTAGTGGTGCTTGACCTTTCC |
| CoiH-R268A-R | GCCAACGGCGGAACGAAAGACGGCAAGTTCCGGAG |
| CoiH-L332A-F | GCTGGCGCCGGACAGGGCCACATTCCGGAACCGCTC |
| CoiH-L332A-R | CTGTCCGGCGCCAGCCGAAGCTACAGCCGCTGAC |

**Table S15. Sequences of *E. coli* codon-optimized LanH genes used in this study.**

| Gene Name | Sequence (5’-3’) |
|---|---|
| CoiH | ATGGATGTGGGCCAGATTGCGAAACGGTTTCCCCTGGTTGCGCGTCCACGCCCTG<br>CGTGTCTGCCACTCGCGGAACGGGTGCGGGAGATTACCGATCTGGCAGCTGCGG<br>CCGAACGCGATGGCAACGTCACCTCAGCAACCGTTGCCCTGAACAAAGCGGCAC<br>TTATCGCCTCTGATTGCGGGATGCCAGATCTCGCTCGCACGTTATGTTGGCGCCATG<br>CCGAGGCAGTGTTACGTGCCCAACCCTTAGGCGCACAAGAAGCGCGTTACGCGTT<br>GGAACCGCTGGTGAATCTGGCGCGCTTGCGTATCCGCGATGGCGACGGACATGGC<br>GCGTATCACCTCTTGGACAGCTTGAACCGTGCCGTTCGCAGCAAAACGGAAGCCG<br>TTATCGATGGGCGCACCACACCGTTGCAGCATCTGACGACATCCGCAGCAGATCAC<br>CAGCAAGTGTGCCAGTGGTTGTGGACCGTGCTGTTAGGTGACGGAACTCGCGCACT<br>GGTTGCGGCAGGCGAATGGGATCGTGCTCGGGCACATAGCCGCCAGCATCGCGG<br>TATTGGTCAACGACTGTTCGATGGTCGCCAGGTTGAGGTACTCGTACGTTGCCTGGG<br>TGAACAGCCGAGTGATGCCCTTATGTTCCTGCATACGAGCCAGCTGGTCGAGCCGT<br>GGGAACAGAGCGTCGCCGCCGCTCTGACAGTCCTGTGTCATCGCGCAGCGGACG<br>AACACCCTGTAGAGCCTGTGGATAAGATGGTGCAGCACTACCTGGCGCTGGACTCT<br>GCTCCGGAACTTGCCGTCTTTCGTTCCCGCGTTGGCTTAGTGGTGCTTGACCTTTCC<br>CCGAAAACCCGCCAAGCGGAGGCGATGCGCCGTCTGACGTGCGAAGCCATGCGT<br>CAGACCGACGGGTATGTTGCCCGTGATGTGCTGGCACATCCGGTCTGTCGTGAAGC<br>CCTGGGTCACGGTGAGCATCGCACTCTGTCAGCGGCTGTAGCTTCGGCTGGCGCC<br>GGACAGGGCCACATTCCGGAACCGCTCACCCAAGCGCTGCTGGGCGCGGCGAG<br>TGTATCGGCCGATGTGTGGGAACGCCACGTCACTCGTCGTGCTAGTCCGGGTTCTT<br>CGCGCAATCTGGACAGCTAA |
| KitH | ATGCATCCGACAGGTCCGATTGCACAGCGCTTTCCGCTGGTGGCGCGGCCACGTC<br>CGGTCTGTCTGCCGCTCGAAGCCCGGGTAGGTCGATTACGGCAACTGGCCGATAC<br>AGCGTCTCAACAATCAAACCCTGGTGAGGCGAGTACTGTGTTTAATCAGGCGGCGTT<br>GTTGGCGAGCGATCTTGGTTTGCCGGACTATGCTCGCGAATTGTGTCATCGACATGC<br>GACGCTTTACCTGACACGGGGACCTCTGCCAGCTATGAGCGCGATACGTTCCCTGG<br>AACCGCTGGTTAATTTAGCCCGCCTCCGTATTCGTGCGGACCATCACCAGCAAGGC<br>CATCAATTGCTGTTGGATCTGTATCGTGCTGTTTCTACATCCACAGAGACCGCGGTTG<br>ACGGCATTCTGGTCCCTGCAAGACTGACTGAAACCGATGAACAGCGTGGCGAGGT<br>GCGCGCTTGGCTGTGGCGCGTGGTGTTAGCTGACGGTACGCGTGCCCTCACATCG<br>GCTGGACGCTGGGACGAGGCACTGCGCCACATTGAAGAGCATCGTGGAATTGGCC<br>AACGTATGTTTGATGGGCGCCAAACCGCGGTGCTGGCCGCGGCAACGAATGGCG<br>ATCTGCCCGCGGCGCGCACTCTCTTAGAATCTACCGAGCCCGGCAGCTCGTGGGA<br>GGATTCTGTTACTGCGGTTCTTACGGCGTTGTGTCGTCCGGGTGATGACAGAGCGGC<br>GGAAACGGCTATCGAACACTGCCTGGCGTTTGAGCCGGATGAGGGGCTTGCAGTTT<br>TCACAATACGTCTGGCTTTAACTGCACTGGATGCCGCATCACTTGACACTCCGGGCA<br>CGCGCAATTTGTTGGCTCAGTTAACTTCCAGAACTGGTGAAACTGGCGATGGTTATGC<br>CCTGCGTGATTTACTGGGGCATGAAGGGGTTCAGCAGCACCTTGCTCCTGCCCAG<br>CGTGAACCTTTAGAACGCGCACTGGCGGCTTGCGCCCTGTCCTCGGGTACCCTTC<br>CTAACGCGTTATCTGGGGTGCTGGAAGAAGCGCTGGCTCATGCTCAAGGCGTGCTT<br>GAAACCTCCCCCTTCGATCAGGGTTCTCCTGGAGGCACTCCACTTGAACAGCGTAA<br>CCGTACCGCGCCCTCTAGCGTAACTTCTACGCCGACACATCCTCTGAATTCCACCC<br>CCTAA |
| LydH | ATGGACGCGTCTCAGATCGTGCGTCGGTTCCCACTGGTAGCCCGTCCGCGTCCCG<br>CATGCAGTCCTCTGGTGGACCGCGTACAGGAGATCGGAGGTTTGGCCCGTGAGGC |

| | |
|---|---|
| | TGAACGTACGGGGCGCCTGGCCCCGGCTTCCGCTGCATTCAACAAAGCTGCACTG<br>CTGGCTTCAGATTGCGCTGTGCCGGATCTTGCTCGTACATTATGTTGGCGTCACGCC<br>GTCGCATATCTGCGGACGCAGCCGTTAGGGGCCCAAGTGGCACGCTTTTCACTGG<br>AACCCTTAGTCAATTTGGCTCGCCTCCTGATCCGCTCTGGTGATGGCGGAGGGGGC<br>TTTGCACTGCTTGATAGTATGTATCGTGCCGTTACGTCAGGCACTGCGGTGACCATAG<br>ATGGGAACGAAGTTAGTTTTGAACGTGTGACTGGAGCCAACGAAGACAGACGTAAAC<br>TGCATCAGTGGCTTTGGACTGTACTGCTGGCGGACGGCACGCGCGCATTAGCTTCG<br>GCCGATATGTGGGCTGAGGCCCATGCACAAGTACAGAGACATAAAGGAATTGGTAGT<br>AGAATGCTGGACGGACGTCAGGTTGCAGTAATTGCCCATATCACAGCAGGAGATATA<br>GAACGTGCTGAGGTTTTACTGTCCGGAACTGTCCCAGGTGAAGCCTGGGAATCGGC<br>AATTGCCGCCCTGCTCACCGTTTTGTGCAGAAGAACGGCGAATTTACCGGTTGATAC<br>CGCATTAGATGATCTGGTAGATCTGTACCGCTCGCTTGAGCGGAGCGCAGGCTTGG<br>CGGTATTTCTGACACGTCTGGGATTATCAGCGATTGACCTCGCCGGTGGCCAAGGG<br>CCTGGGATCGCATCGCGTATCGCCACGGATCTGGTTGAAGAAACCGTGGCGGTAG<br>GCGATGGATACGCAGCTCGCGATGTACTGAGTCATCCAGGATGCCATGAATATATGA<br>CGGCCCAACAGGCTGATCGTTTGCAGCGGATTGTCGAAGGATGTGCTTTGGGTTCG<br>GGTGCGATTCCGGAAGACCTGAGAGGGGACTTATTTACGGCCTTGAACACCAGTGA<br>GAGTGTCATCACGAGAGTGTGCGCTGCGGCGCCATCAGCTTCCGCATAA |
| PauH | ATGGTCGATCTCGATCGTATTGCCCGTCGTTTTCCATTGGTCGCCCGTCCGCGACCA<br>GCCTGTCTCGCGCTGGAAGAGCGGGTAAAACAGGTTGGGGAATTAGCACGCGCC<br>GCGCGGGCAGGTGAAGGTGATCGGCTCGCCCGCGCTGCGAGCGCCTCGAACTT<br>GGCGGCATTAATTGCTAGTGATTGTGGAATGCCGGATCTGGCACGTACACTGTGTTG<br>GCGTCAATTTGATGTATATATGCGCGCGAGACCTTTGGACGCACCCTCTGCGCGTCA<br>TGCGCTGGAGCCAATTGTGAATTTAGCACGGCTCCATATCCGCGACGGTGATGGTG<br>CATCGGCCTATTCGCTTTTAGATACGTTATATCGGGCCGTGCGGTCAGCAACGAAAG<br>GCCTGATAGATGGCCGTCAAGTTTCTTTTCAAGGCTTCACAGCATCCGCCGAAGATC<br>ATAGAACCCTCTGCCAGTGGCTTTGGACTGTTGTCCTGGCTGAAGGGACACGCGCG<br>CTGGCAACCGCGGGTAGATGGAGTGAAGCGCTGGCCCATGCGGAGCAGCATCGA<br>GGCGTTGGAGATCGTTTAATGGATGGCCGTCAAGTGGCTATACTCGCCAACTGCTTG<br>GCCGGTGATCCTGCTGCGGCACTGGCAATTCTCGAAGCGTCGGCTATTAGCGAAC<br>CCTGGGAGGGCGCAGTAGCCGCATGTCTGAAAGTGTTCTGCTTGAATGCATCAGCC<br>CGTCCCGTGGGCGCGGCCGTCGATGAAATGGTCGAGCGTTACCTTGCACTTGATAC<br>GGCGCCCGAGTTAGTTGTATTCCGCACGCGCGTGGGTCTGTCCTTAGTTGATTTGGC<br>GGATGGAGCAGACCAGGCAGACGTGTCCCAAGCGACCACGCGCCTGCTGGATGA<br>AGCAGTTACCGCTAGAGATGGATATGCAGGTCGCGAGGTGCTGGCACATAACGCGT<br>CTCGCGCGCGTCTCACGCTCACCCAAGAACGCATACTGCGTGCGGTTGTTGAGTC<br>GTCTCGTTTAGGTCGCGGGATAATGTCAGCCCAAATGGTCGCAGATCTGTTAAAGGC<br>TATGTCTATGAGCGAGGCCGCGACGGTATAA |

**References**

(1) Nguyen, D. T.; Zhu, L.; Gray, D. L.; Woods, T. J.; Padhi, C.; Flatt, K. M.; Mitchell, D. A.; van der Donk, W. A. Biosynthesis of macrocyclic peptides with C-terminal β-amino-α-keto acid groups by three different metalloenzymes. *ACS Cent. Sci.* **2024**, *10*, 1022-32.
(2) Brademan, D. R.; Riley, N. M.; Kwiecien, N. W.; Coon, J. J. Interactive peptide spectral annotator: A versatile web-based tool for proteomic applications. *Mol. Cell. Proteom.* **2019**, *18*, S193-S201.
(3) Tietz, J. I.; Schwalen, C. J.; Patel, P. S.; Maxson, T.; Blair, P. M.; Tai, H. C.; Zakai, U. I.; Mitchell, D. A. A new genome-mining tool redefines the lasso peptide biosynthetic landscape. *Nat. Chem. Biol.* **2017**, *13*, 470-8.
(4) van Kempen, M.; Kim, S. S.; Tumescheit, C.; Mirdita, M.; Lee, J.; Gilchrist, C. L. M.; Söding, J.; Steinegger, M. Fast and accurate protein structure search with Foldseek. *Nat. Biotechnol.* **2024**, *42*, 243-6.
(5) Holm, L.; Laiho, A.; Törönen, P.; Salgado, M. DALI shines a light on remote homologs: One hundred discoveries. *Protein Sci.* **2023**, *32*, e4519.
(6) Oberg, N.; Zallot, R.; Gerlt, J. A. EFI-EST, EFI-GNT, and EFI-CGFP: Enzyme Function Initiative (EFI) web resource for genomic enzymology tools. *J. Mol. Biol.* **2023**, *435*, 168018.
(7) Katoh, K.; Rozewicki, J.; Yamada, K. D. MAFFT online service: multiple sequence alignment, interactive sequence choice and visualization. *Brief Bioinform.* **2019**, *20*, 1160-6.
(8) Polak, L.; Skoda, P.; Riedlova, K.; Krivak, R.; Novotny, M.; Hoksza, D. PrankWeb 4: a modular web server for protein-ligand binding site prediction and downstream analysis. *Nucleic Acids Res.* **2025**, *53*, W466-W71.
(9) Shi, Y.; Bueno, A.; van der Donk, W. A. Heterologous production of the lantibiotic Ala(0)actagardine in *Escherichia coli*. *Chem. Commun.* **2012**, *48*, 10966-8.
(10) Ijichi, S.; Hoshino, S.; Asamizu, S.; Onaka, H. SolS-catalyzed sulfoxidation of labionin to solabionin drives antibacterial activity of solabiomycins. *Bioorg. Med. Chem. Lett.* **2023**, *89*, 129323.
(11) Ye, Y.; Minami, A.; Igarashi, Y.; Izumikawa, M.; Umemura, M.; Nagano, N.; Machida, M.; Kawahara, T.; Shin-Ya, K.; Gomi, K.; Oikawa, H. Unveiling the biosynthetic pathway of the ribosomally synthesized and post-translationally modified peptide ustiloxin B in filamentous fungi. *Angew. Chem. Int. Ed.* **2016**, *55*, 8072-5.
(12) Seebeck, F. P. In vitro reconstitution of Mycobacterial ergothioneine biosynthesis. *J. Am. Chem. Soc* **2010**, *132*, 6632-3.
(13) Braunshausen, A.; Seebeck, F. P. Identification and characterization of the first ovothiol biosynthetic enzyme. *J. Am. Chem. Soc.* **2011**, *133*, 1757-9.
(14) Sarksian, R.; van der Donk, W. A. Divergent evolution of lanthipeptide stereochemistry. *ACS Chem. Biol.* **2022**, *17*, 2551-8.
